# Ancestry-restricted expression of P2RX5 defines a CAR T-cell target across diverse blood cancers

**DOI:** 10.64898/2026.01.14.698748

**Authors:** Zhiwei Ang, Annette Castro, Luca Paruzzo, Carolin Schmidt, Tori J. Fuller, Alberto Carturan, Katharina E. Hayer, Christopher Kwok, Jacinta Davis, Samantha S. Soldan, Federico Stella, Manuel Torres-Diz, Rawan Shraim, Patricia King Sainos, Kayla Ji, Patrick J. Krohl, Justyn Fine, Shannon E. McGettigan, Priyanka Sehgal, Daniel Martinez, Patrizia Porazzi, Jamie B. Spangler, Paul M. Lieberman, James L. Riley, Vinodh Pillai, Haley Newman, Martin Carroll, Zainul S. Hasanali, David Allman, Dan T. Vogl, Caroline Diorio, David T Teachey, Marco Ruella, Andrei Thomas-Tikhonenko

**Affiliations:** Division of Cancer Pathobiology, Children’s Hospital of Philadelphia, Philadelphia, PA; Center for Cellular Immunotherapies, Perelman School of Medicine, University of Pennsylvania, Philadelphia, PA; Division of Hematology/Oncology, Perelman School of Medicine, University of Pennsylvania, Philadelphia, PA; Division of Oncology, Children’s Hospital of Philadelphia, Philadelphia, PA; Department of Biomedical and Health Informatics, Children’s Hospital of Philadelphia, Philadelphia, PA; Vagelos Life Sciences Management Program, University of Pennsylvania, Philadelphia, PA, USA; Genome Regulation and Cell Signaling Program, The Wistar Institute, Philadelphia, PA; Department of Biology, College of Arts and Sciences, University of Pennsylvania, Philadelphia, PA; Department of Chemical & Biomolecular Engineering, Johns Hopkins University School of Engineering, Baltimore, MD; Department of Biomedical Engineering and Translational Tissue Engineering Center, Johns Hopkins University School of Medicine, Baltimore, MD; Program in Molecular Biophysics, Johns Hopkins University, Baltimore, MD; Department of Pathology and Laboratory Medicine, Perelman School of Medicine, University of Pennsylvania, Philadelphia, PA; Pathology Core, Children’s Hospital of Philadelphia, Philadelphia, PA; Department of Microbiology, Perelman School of Medicine, University of Pennsylvania, Philadelphia, PA; Division of Hematopathology, Children’s Hospital of Philadelphia, Philadelphia, PA; Department of Pediatrics, Perelman School of Medicine, University of Pennsylvania, Philadelphia, PA

## Abstract

Purinergic receptor P2RX5 can serve as an HLA-restricted graft-versus-leukemia antigen ^1^. Yet, its direct targeting with chimeric antigen receptor (CAR) T cells remains unexplored, due to the prevailing view that protein-truncating mutations have rendered human *P2RX5* a pseudogene ^2^. Here, leveraging transcriptomic datasets and newly-developed monoclonal antibodies, we show that up to 75% of individuals with African ancestry express the full-length variant of P2RX5 (P2RX5-FL). While in healthy tissues *P2RX5* is transcribed only in B cells and prenatal thymocytes, it is upregulated across multiple hematologic cancers including B-cell lymphomas (BCL), multiple myelomas (MM), and T-cell acute lymphoblastic leukemias (T-ALL). Crucially, unlike its truncated counterparts, P2RX5-FL protein is stable and plasma membrane-bound. We reformatted one monoclonal antibody as a P2RX5-directed CAR and expressed it in donor T cells. These CART-X5 cells were not prone to fratricide and were effective in reducing disease burden and extending survival in xenograft mouse models of T-ALL and MM. Together, these findings nominate P2RX5-FL as an immunotherapy target for BCL and MM patients that have relapsed after CD19 or BCMA-directed therapies with antigen-negative disease^3–6^. P2RX5-FL could also be targeted in chemoresistant T-ALL, where new immunotherapies avoiding CAR T-cell fratricide and T-cell aplasia are urgently needed ^7,8^.

## Individuals with African ancestry express a stable, full-length variant of *P2RX5* (*P2RX5-FL*)

Protein-truncating mutations are known to contribute to P2RX5 pseudogenization ^9^, but their prevalence across human populations remains incompletely understood. Analysis of 1000 Genomes haplotype data ^10^ reveals two common variants: a frameshift caused by a single-nucleotide deletion in exon 3 (rs3215407 delG) generating a C-terminally truncated variant (ΔC) and an intron 10 splice-site substitution (rs891746 A→C) associated with exon 10 skipping (Δ10) (**Fig. 1a**). Yet expression of the full-length (FL) protein requires preservation of both ancestral alleles: rs3215407 G and rs891746 A, which per LDpair ^11^ are tightly linked. Of note, the productive “(G)5 + A” is more prevalent in African ancestry (AF) populations than in the pooled, global population of the 1000 Genomes dataset (66% versus 18%). Assuming Hardy-Weinberg equilibrium, the probability of inheriting at least one copy of the “(G)5 + A” haplotype is 0.33 for all ancestries and 0.89 for AF individuals. The latter might be an overestimation, since in larger AF cohorts, the combined frequency of rs891746 A homozygous (A/A) and heterozygous (A/C) carriers doesn’t exceed 75% (**Fig. 1b**). Still, the majority of these donors are expected to harbor the far more common “(G)5 + A” haplotype (66% frequency) and not the rare “delG + A” haplotype (2% frequency) (**Fig. 1a**), and thus encode P2RX5-FL.

While the rs891746 SNP genotype is associated with *P2RX5* exon 10 skipping based on genotyping and RNA analyses ^9^, we set out to establish a causal relationship. Upon transfection into Raji and OCI-ly8 B-cell lines, *P2RX5* minigenes carrying the rs891746 C allele yielded exon 10-skipping transcripts, while those with the A-allele produced longer, exon 10-including transcripts (**Extended Data Fig. 1a**). We also used the CRISPR-Cas9-based homologous recombination system to edit the endogenous *P2RX5* gene in DoHH2 cells, successfully converting 3% of its homozygous rs891746 C alleles (C/C) to A (C→A) (**Fig. 1c**, top panel). We then designed qPCR primers that distinguished exon 10 inclusion from skipping on control DNA templates, alongside a primer set targeting constitutive exon 9 (**Extended Data Fig. 1b**). Our RT-qPCR assay detected an 8-fold increase in the exon 10 inclusion signal for the C→A-edited cultures, relative to C→C and parental controls (**Fig. 1c**, bottom panel).

The inclusion of *P2RX5* exon 10 is consequential. After transient transfection into adherent, non-lymphoid cells, the FL protein was reported to colocalize with the cell membrane while the Δ10 variant was predominantly cytoplasmic ^9^. To analyze subcellular localization of P2RX5 variants in B-lymphoid cells, we inserted the FLAG tag into the predicted extracellular loop of P2RX5 (**Fig. 1d**) and transfected these constructs into Raji and Ramos cells. While Δ10-FLAG and FL-FLAG were expressed equally well at the mRNA level (**Extended Data Fig. 1c**, top panel), only FL-FLAG was detected on live, non-permeabilized cells via flow cytometry (**Extended Data Fig. 1c,** bottom panel). We also knocked the FLAG tag sequence into the endogenous P2RX5 locus in Raji cells, an endemic Burkitt lymphoma cell line of African origin with the homozygous *P2RX5-FL* haplotype. After confirming the insertion via RT-PCR and amplicon re-sequencing (**Fig. 1e**, top panel), we readily detected FL-FLAG expression on the cell surface using flow cytometry, where P2RX5 KO cells served as a negative control (**Fig. 1e**, bottom panel**; Extended Data Fig. 1d)**. In addition to disrupting membrane localization, exon 10 exclusion may also compromise protein stability by removing the predicted second transmembrane helix of P2RX5 (**Fig. 1d**). To test this, we ectopically expressed FL-GFP and Δ10-GFP fusion proteins in Raji cells. Upon cycloheximide (CHX) treatment to halt new protein synthesis, flow cytometry at different time points revealed that Δ10-GFP signal diminished more rapidly that the FL-GFP signal (**Fig. 1f**). This suggests that, at least in the Raji B-lymphoma line, P2RX5-Δ10 is rapidly degraded while P2RX5-FL is stably expressed on the cell surface and is potentially targetable with CAR-based immunotherapies.

## BCL, MM, and T-ALL neoplasms upregulate P2RX5 mRNA

The range of *P2RX5* expression across human blood cancers is not fully elucidated. Using aggregate microarray data from the HEMAP portal ^12^, we observed elevated *P2RX5* mRNA levels not only in B-cell acute lymphoblastic leukemias (B-ALL), B-cell lymphomas (BCL) and chronic lymphocytic leukemias (CLL)—but also, unexpectedly, in MM and T-ALL, especially within two high-risk T-ALL subtypes, LMO2 γδ-like and ETP-like (**Fig. 2a and Extended Data Fig. 1e**). Additionally, high levels of P2RX5 expression were apparent in the RNA-seq datasets corresponding to MM samples from the MMRF CoMMpass study ^13^ and endemic Burkitt’s lymphoma (eBL) cohorts from West Kenya ^14^ and Uganda ^15,16^, including ∼75% of samples with the FL haplotype (**Extended Data Fig. 1f**). This enhanced P2RX5 expression was retained by MM, T-ALL, BCL and CLL cell lines from the Cancer Cell Line Encyclopedia (CCLE) (**Fig. 2b**). Ranking high among them were two BCL (Daudi, Raji) and two MMs (MM.1s, RPMI-8226) cell lines with the FL/FL-genotype, as well as a T-ALL cell line (MOLT-16) with the FL/ΔC genotype (**Fig. 2b, c**). The expression of exon 10-containing *P2RX5-FL* mRNA by these cell lines was confirmed via short-read RNA-seq (**Fig. 2c**), RT-qPCR, and long-read direct RNA-seq ^17^ (**Extended Data Fig. 1g, h**).

## P2RX5-FL protein is expressed in blood cancer cell lines and patient samples

We next developed two monoclonal antibodies, 1D9 and 2F4, which could recognize P2RX5-FL protein in live-cell flow cytometry assays. Both antibodies stained Raji and MM.1s cells with native P2RX5-FL expression and Ramos-P2RX5-FL transfectants (**Fig. 3a**). 1D9 and 2F4 had binding affinities in the 43-71-nM range (**Extended Data Fig. 2a**) and recognized non-identical epitopes, as revealed in competition assays (**Extended Data Fig. 2b**). In immunoblots, 1D9 detected a ∼60 kDa band in all *P2RX5-FL* haplotype cell lines (**Fig. 3b, c**); its intensity was increased in Raji cells overexpressing P2RX5-FL but not P2RX5-Δ10 mRNA (**Extended Data Fig. 2c**), consistent with the rapid degradation of Δ10 protein (**Fig. 1f**). Similarly, Δ10/Δ10 DoHH2 cells didn’t yield the 1D9-reactive band, but their rs891746 C→A-edited, FL-encoding derivatives did (**Fig. 3b**); this result was confirmed in flow cytometry experiments **(Extended Data Fig. 2d**). Enzymatic deglycosylation or two N-to-A substitutions (N77A/N202A) shifted the apparent molecular weight of the 1D9-reactive protein band from ∼60 to ∼50 kDa (**Extended Data Fig. 2e**), which is in line with the 49 kDa predicted molecular weight of P2RX5-FL polypeptide.

We further quantified 1D9 binding to cell-surface P2RX5 in various cell lines using antibody-binding capacity (ABC) assays (**Fig. 3d**) and found it to be concordant with mRNA levels (**Fig. 2c**). The densities of the 1D9-accessible P2RX5 were estimated at 3,000-6,000 molecules/cell for cell lines with the homozygous FL/FL haplotype and as ∼1,000 molecules/cell in the FL/ΔC MOLT-16 cells (**Fig. 3d**). While these values were substantially lower than the 40,000-100,000 CD19 molecules/cell, they were comparable to ∼6,000 molecules/cell estimated for B-cell maturation antigen (BCMA) in MM.1s and RPMI-8226 cells, which are known to be sensitive to BCMA-directed CAR T cells ^18^. Finally, we determined that P2RX5-FL expression persists in these cell lines after exposure to common chemotherapy drugs (i.e., doxorubicin, vincristine, bortezomib, lenalidomide and dexamethasone), although it was reduced in T-ALL cell lines upon treatment with DAPT, a γ-secretase/Notch inhibitor, a class of drugs in clinical development for T-ALL ^19,20^ (**Extended Data Fig. 3a-e**). Furthermore, P2RX5 expression was unaffected by the knockout of genes encoding other targets for immunotherapy, namely CD19, CD20, CD22, and CD79B in BCL lines and CD38 and BCMA in MM cell lines; nor was their expression compromised by the loss of P2RX5 (**Extended Data Fig. 3f-h**).

To extend these findings to cancer patient samples, we selected formalin-fixed, paraffin-embedded (FFPE) biopsies of human BCL and MM lesions, as well as viably frozen bone marrow aspirates from MM and T-ALL patients, and splenic cells from T-ALL xenograft-bearing mice. All samples were subject to Taqman genotyping assays to determine their P2RX5 haplotypes. FFPE sections were stained with the 1D9 antibody using immunohistochemistry (IHC), while live cell suspensions were stained with both 1D9 and 2F4 in flow cytometry assays. These analyses revealed robust, FL haplotype-dependent expression of P2RX5 in BCL and MM lesions (where it colocalized with CD20 and CD138, respectively) (**Fig. 4a, b; Extended Data Fig. 4**), in CD138-positive bone marrow mononuclear cells (BMMNC) from adult MM cases, in abnormal BMMNCs from pediatric T-ALL cases (**Fig. 4c**), and in pediatric T-ALL PDXs (**Fig. 4d; Extended Data Fig. 5**).

In order to establish the specificity of staining with 1D9 and 2F4, we confirmed lack of cross-reactivity with other P2RX family members (**Extended Data Fig. 6a**). The 1D9 binder was also screened against over 6,000 human transmembrane proteins represented on the Integral Molecular Membrane Proteome Array ^21^ (**Extended Data Fig. 6b**). This screen revealed robust binding of 1D9 to P2RX5-FL and weak, off-target binding to Nectin-4, which was only evident in ectopic overexpression models (HEK293T and RPMI-8402 cells) but not on carcinoma cell lines with endogenous Nectin-4 overexpression (T-47D and HT-1376 cells) (**Extended Data Fig. 6c, d**).

## P2RX5FL-directed CAR T cells exhibit potent antitumor activity while avoiding fratricide

Given high specificity of 1D9 binding, we developed a P2RX5-directed, 1D9 scFv-based second-generation CAR featuring 4-1BB costimulatory and CD3ζ signaling domains (**Fig. 5a**). This 1D9-BBz CAR or CART-X5 construct was cloned into a lentiviral vector and transduced into T cells from several independent donors, including 3 *FL*/*FL*-genotyped donors of AF ancestry (**Extended Data Fig. 6e**). Compared to untransduced controls, 1D9-BBz had increased avidity towards P2RX5FL-positive RPMI-8226 cells (**Fig. 5b**) as well as measurable interferon γ (IFNG) secretion upon exposure to P2RX5FL-positive MM.1S cells (**Fig. 5c**). Importantly, IFNG secretion by 1D9-BBz remained at baseline when they were challenged with P2RX5FL-negative controls, including Nectin-4-expressing keratinocytes and T-47D and HT-1376 cells (**Fig. 5c**). Of note, 1D9-BBz T cells were comparable to untransduced controls in terms of *in vitro* expansion rates, exhaustion marker expression (LAG-3, TIM-3, PD-1) as well as differentiation into effector and memory subsets (**Extended Data Fig. 6f-h**).

To evaluate its cytolytic activity, 1D9-BBz T cells were co-cultured with target cell lines expressing luciferase (luc+) at a 3:2 effector-to-target (E:T) ratio. After one day, luc+ target cell viability was quantified by bioluminescence and normalized to target-alone controls. Compared to control T cells, 1D9-BBz T cells eliminated at least 50% of cells that expressed P2RX5-FL either endogenously (RPMI-8226, MM.1s, Raji) or ectopically (Ramos-P2RX5FL, RPMI-8402-P2RX5FL). Crucially, this cytolytic activity was maintained against engineered models of antigen-escape (MM.1S-BCMA KO and Raji-CD19 KO), but negligible against P2RX5-FL-negative target cells (Ramos-Puro, RPMI-8402, RPMI-8402-NECTIN4, MM.1S-P2RX5KO, Raji-P2RX5KO) (**Fig. 5d**). Even at a low E:T ratio of 1:16, 1D9-BBz T cells controlled MOLT-16 and MM.1S target cell outgrowth over 4 days (**Extended Data Fig. 6i**). When MM.1s cells were pretreated with 1μM dexamethasone, a frontline MM therapeutic, their susceptibility to 1D9-BBz T cells was enhanced further (**Extended Data Fig. 6j, k**). This was in line with our observation that dexamethasone upregulates cell-surface P2RX5 in MM.1s (**Extended Data Fig. 3c, d**), and a prior report that 1μM dexamethasone does not impair CAR T-cell cytolytic activity ^22^. The cytolytic activity of 1D9-BBz T cells was also durable. When initial cultures of 4,000 1D9-BBz T cells were challenged every 3 days with 8,000 additional MM.1s target cells, flow cytometric monitoring revealed that 1D9-BBz T cells suppressed MM.1s outgrowth for at least four sequential challenge rounds, while control T cells displayed little inhibitory activity (**Fig. 5e**).

There is evidence in the literature and in public RNA-seq datasets that *P2RX5* mRNA is transiently induced during *ex vivo* activation of normal ^23^ and CD19-directed CAR T cells ^23,24^, although we found no evidence for its expression *in vivo*, even during peak expansion CART-19 cells in infused patients ^25^ (**Extended Data Fig. 7a-c**). Nevertheless, we used 1D9 and 2F4 to measure cell-surface P2RX5 levels on *ex vivo* activated donor T cells during 1D9-BBz CAR manufacturing. While we detected no increase in P2RX5 immunostaining for *Δ10/ΔC*-genotyped T cells, we did detect a modest increase for *FL/FL* and *FL/Δ10* T cells (**Extended Data Fig. 7d-f**). This increase was short-lived, peaking between days 5 and 7 post-activation before returning to baseline by day 12. To assess the functional significance of transient P2RX5 expression, we compared *FL/FL* 1D9-BBz T cells to those from P2RX5-FL-negative, *Δ10/ΔC* donors. We found no meaningful difference in their expansion, exhaustion marker expression, effector memory differentiation, and P2RX5FL-specific cytotoxicity in vitro (**Fig. 5e; Extended Data Fig. 6f-j**). Furthermore, activated T cells expansion was not suppressed during coculture with autologous 1D9-BBz T cells from the same *FL/FL* donor (**Extended Data Fig. 7g**). Taken together, these findings ruled out sustained patterns of 1D9-BBz CAR-mediated *FL/FL* T-cell fratricide.

To test their *in vivo* potency, we intravenously administered 8×10^5^ 1D9-BBz CAR T cells (*Δ10*/*ΔC*) into mice bearing bone marrow xenografts of human RPMI-8226 MM cells (*FL*/*FL*). A lower dose of 300,000 CAR T cells (*FL*/*FL*) per mouse, was administered into two T-ALL PDX models (ETP-8, *FL*/*FL* and ALL-8, *FL*/*ΔC*). All xenografted MM and T-ALL cells expressed luciferase, which allowed for longitudinal tracking of tumor burden via whole-animal bioluminescent imaging (**Fig. 5f-h**). Relative to mice treated with donor-matched control T cells, 1D9-BBz CAR treatment induced tumor regression in RPMI-8226 models, and blunted tumor progression in ETP-8 and ALL-8 models relative to donor-matched controls (**Fig. 5g**, top panels). Furthermore, 1D9-BBz CAR treatment extended survival for the RPMI-8226 (>100 days) and ETP-8 ( >50 days) models (**Fig. 5g**, bottom panels). In the RPMI-8226 model, 1D9-BBz CAR T cells persisted for over 3 months and achieved therapeutic outcomes comparable to those achieved with an equivalent dose of the clinical BCMA CAR, ciltacabtagene autoleucel (**Fig. 5i**).

## P2RX5 is expressed by prenatal thymocytes and B cells, but is B-cell restricted in postnatal tissues

To assess potential CART-X5 on-target/off-tumor toxicities, we took advantage of flow-sorted immune subsets across bone marrow and tonsils samples characterized by us earlier ^26,27^ and published datasets of peripheral blood mononuclear cells ^28,29^ (**Fig. 6a, b; Extended Data Fig. 8a**). We found that P2RX5 expression is undetectable across most hematopoietic lineages including CD34^+^ hematopoietic stem, progenitor cells and T-lymphoid cells, but it is strongly induced in B-lymphoid cells beginning at the pre-B and immature B-cell stage. Further analysis of bulk RNA-seq data from the Genotype-Tissue Expression (GTEx) ^30,31^ and Human Protein Atlas cohorts ^28^ showed the highest *P2RX5* expression levels in B cell-rich lymph node, tonsil, spleen, appendix and small intestine, with little to no expression across other organs (**Extended Data Fig. 8b-d**).

**Fig. 1.**
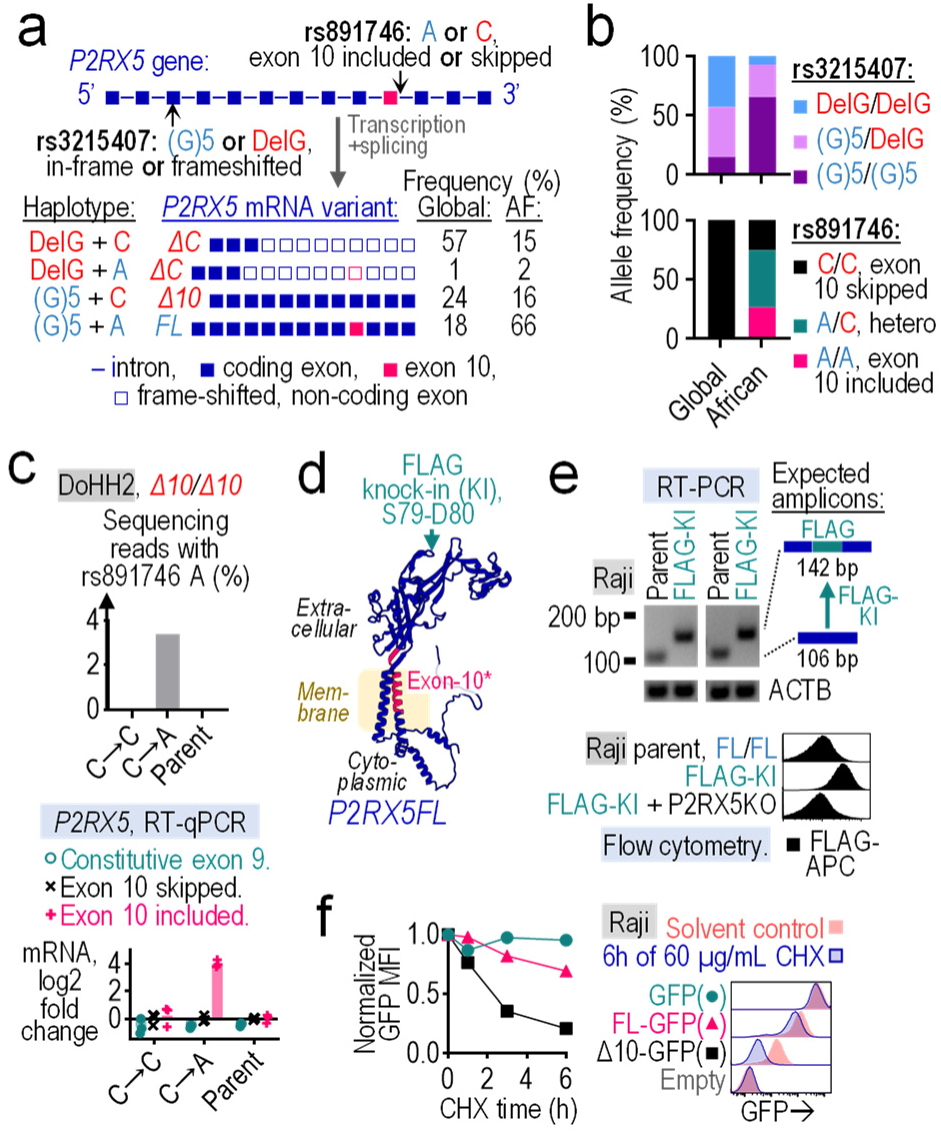
Individuals of African ancestry express the full-length cell surface variant of P2RX5 (P2RX5-FL). **a.** Top: Exon-intron structure of human *P2RX5* with the locations of the rs3215407 indel and the rs891746 SNP. Bottom: *P2RX5* variants that arise due to rs3215407 and rs891746, and their population haplotype frequencies from the NCBI LDlink 7.3.0 release. AF, African. **b.** Aggregate allele frequencies of rs3215407 and rs891746, from the NCBI SNP database (ALFA project release 4). **c.** Analysis of P2RX5-Δ10/Δ10-genotyped DoHH2 cells before (parent) and after CRISPR-Cas9-mediated genome editing of rs891746 from C-to-C (C→C) or from C-to-A (C→A). Top: Percent of Illumina targeted sequencing reads with rs891746 A. Bottom: *P2RX5* mRNA fold change relative to unedited parent cells. RT-qPCR primers measure constitutive exon 9, as well as the skipping or inclusion of exon 10. **d.** Ribbon diagram of P2RX5-FL protein structure, as predicted by AlphaFold 2, with the FLAG tag knock-in (FLAG-KI) site. **e.** Analysis of unmodified Raji parent cells, a *P2RX5* FLAG tag knock-in (FLAG-KI) daughter line, and the FLAG-KI followed by *P2RX5* knockout (P2RX5KO) double-modified line. Top: RT-PCR amplification of the region flanking the FLAG insertion site on *P2RX5* mRNA. Amplicons were separated by agarose gel electrophoresis. Bottom: Live-cell flow cytometry of cell-surface FLAG. **f.** Live-cell flow cytometry of P2RX5(Δ10 or FL)-GFP fusion protein abundance in Raji cells at various time points following cycloheximide (CHX) treatment.

**Fig. 2.**
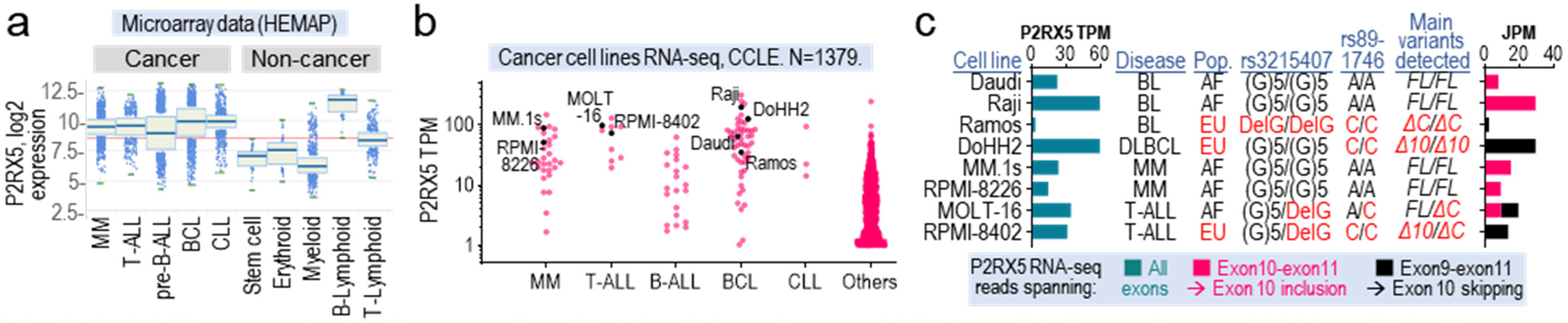
P2RX5 transcription is upregulated in B-cell lymphomas (BCL), multiple myelomas (MM) and T-cell acute lymphoblastic leukemia (T-ALL). **a.** Collated microarray data of P2RX5 expression in ∼10,000 normal and neoplastic leukocyte samples across all lineages, as published on the HEMAP portal. **b, c.** RNA-seq analysis of *P2RX5* transcript abundance (Transcripts Per Million, TPM) across all cell lines in the Cancer Cell Line Encyclopedia (CCLE, Broad 2019) (**b**), and in a selection of cell lines (**c**). Quantitation of RNA-seq reads corresponding to exon 10 skipping (exon 9-exon 11 junction) and inclusion (exon 9-exon 10 junction). JPM: junction reads per million. The rs3215407 and rs891746 alleles were determined using Sanger sequencing. AF, African. EU, European.

**Fig. 3.**
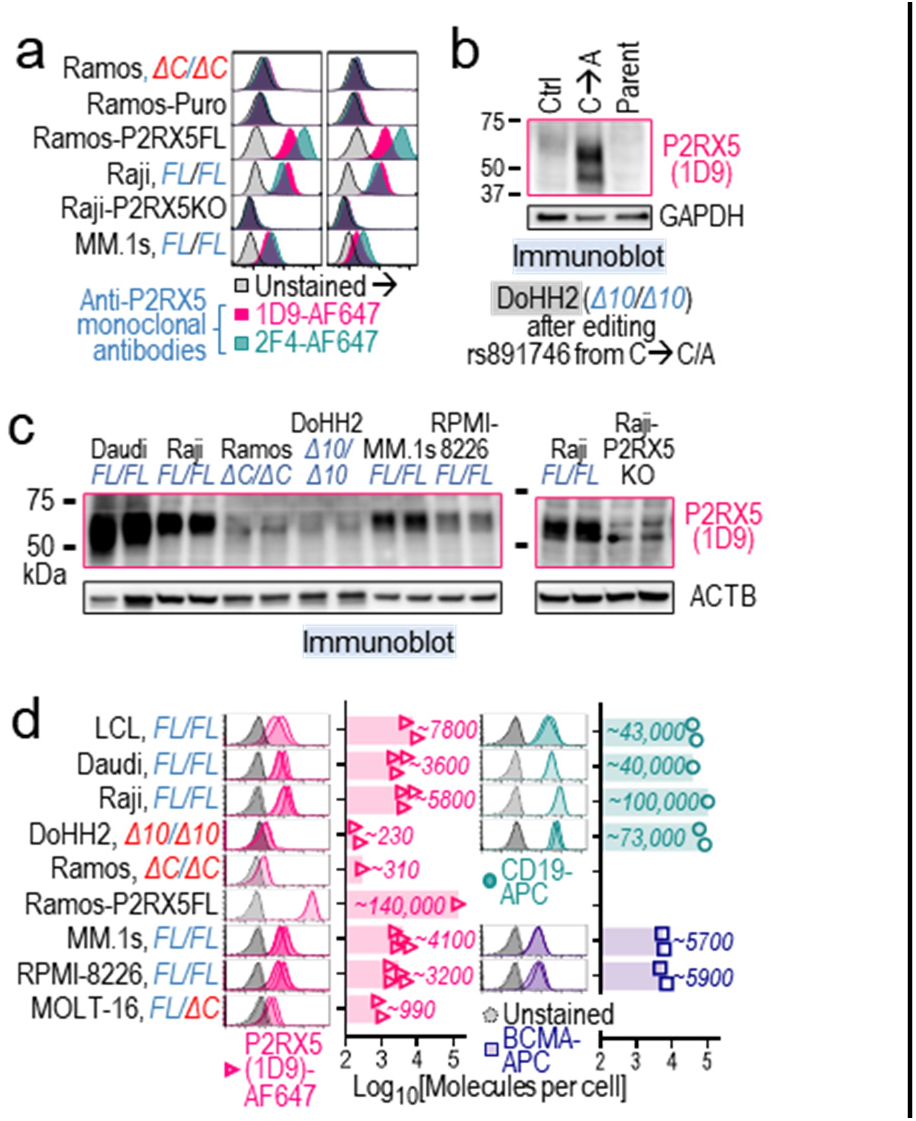
Elevated P2RX5 protein in cell lines expressing the *FL* variant. **a.** Live-cell flow cytometry of Ramos, Raji and MM.1s cell lines after staining with unlabeled P2RX5-reactive monoclonal antibodies (1D9 or 2F4), followed by indirect staining with an Alexa Fluor 647(AF647)-labeled secondary antibody. Ramos-P2RX5 FL cells stably overexpress the full length protein. **b, c**. Representative immunoblots showing P2RX5 expression in cell lines, as detected via 1D9 antibody. Panel b shows DoHH2 cell before (parent) and after CRISPR-Cas9-mediated genome editing of rs891746 from C-to-C (C→C) or from C-to-A (C→A). Panel c shows unedited parental cell lines and a Raji P2RX5 knockout (P2RX5 KO). **d.** Live-cell flow cytometry of cell lines stained with the indicated antibodies. Bar graphs show the average number of antibody molecules bound per cell, as determined via Antibody Binding Capacity (ABC) assays. Each datapoint represents an independent experiment.

**Fig. 4.**
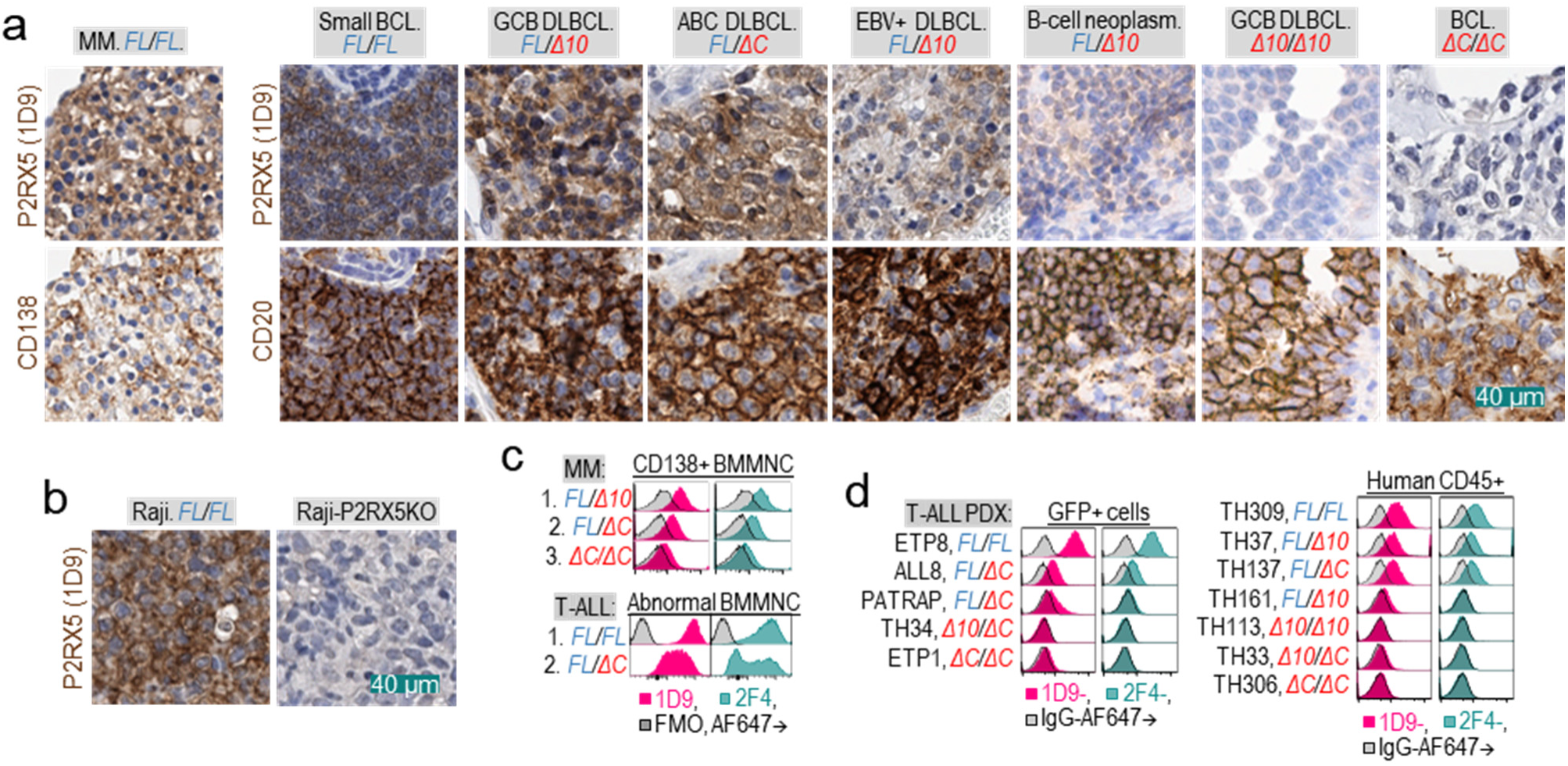
Patient tumors with the FL haplotype display elevated P2RX5 protein levels. **a.** Representative IHC images of tumor lesions from patients with MM and BCL. FFPE sections were stained as indicated for P2RX5 (via 1D9 antibody), CD138, or CD20 using DAB substrate (brown), and with hematoxylin nuclear counterstaining (blue). Refer to Extended Data Fig. 4 for images of the whole tumor sections. **b.** IHC images for mice-engrafted Raji parent and P2RX5 knockout (P2RX5KO) tumors after immunostaining with 1D9. **c, d.** Flow cytometry of bone marrow mononuclear cells (BMMNCs) from patients with MM or T-ALL (**c**), and of splenic cells from T-ALL PDX models (**d**). Histograms display staining intensities for IgG isotype control, 1D9, or 2F4 antibodies within indicated population gates (CD138+, GFP+, and human CD45+). GFP or human CD45 expression distinguish human T-ALL xenografts from murine cells. For the bottom panel of **c**, the abnormal T-lymphoblast population is defined as CD14-CD45+cCD3+ CD7+ CD5+CD4-CD8-CD34-, based on clinical pathology reports. FMO, fluorescence minus one control. Flow cytometry gating examples for panels c to d are shown in Extended Data Fig. 5.

**Fig. 5.**
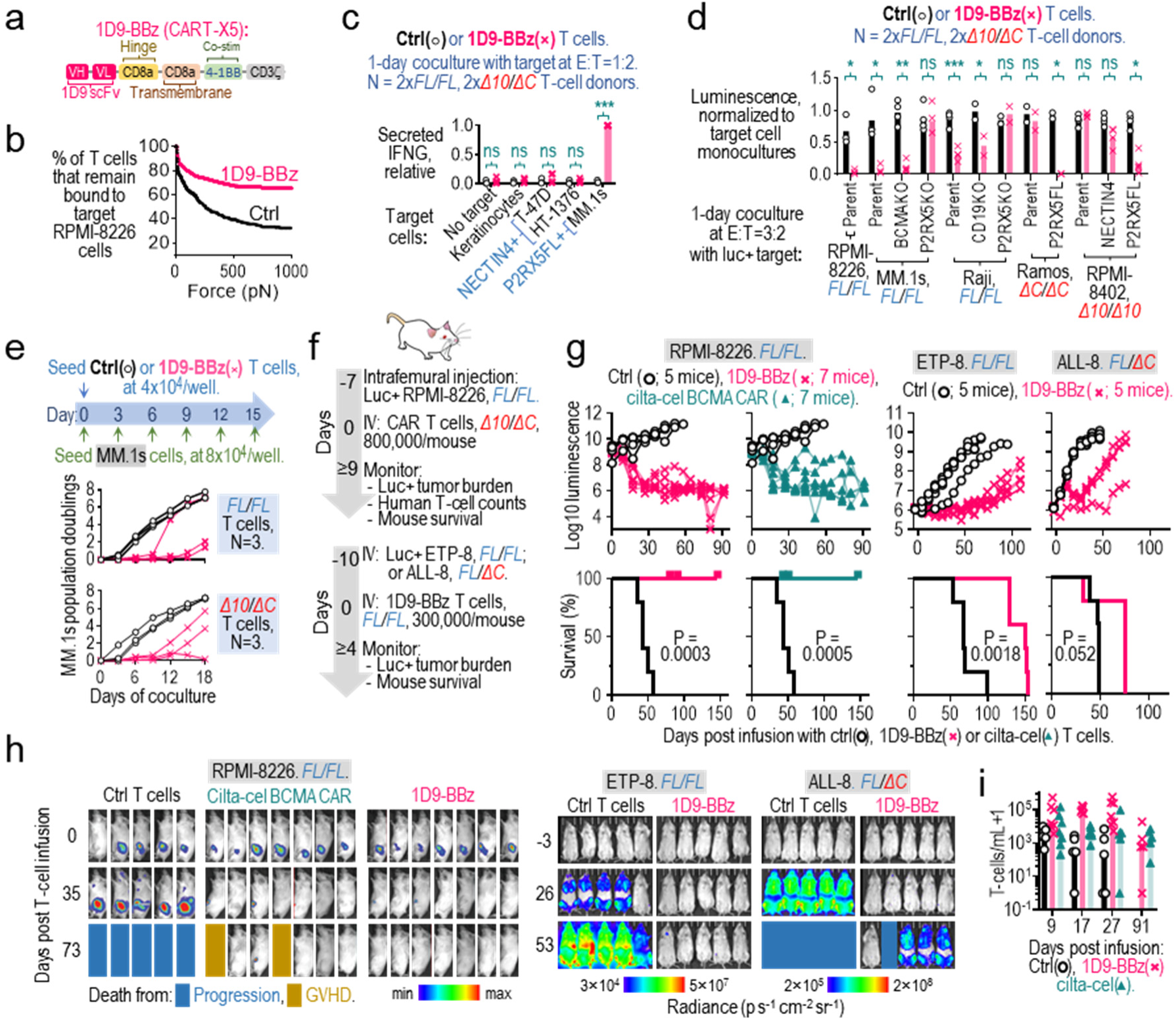
1D9-BBz CAR T cells (CART-X5) display potent anti-tumor activity against *P2RX5 FL*-positive tumors. **a.** Design of the P2RX5-directed second-generation CAR, 1D9-BBz. **b–e.** Functional characterization of 1D9-BBz T cells relative to untransduced controls during coculture with the indicated target cells at specified effector-to-target (E:T) ratios. Assayed functions include: **b**, cell-to-cell avidity (via LUMICK z-Movi instrument); **c**, IFNG secretion, normalized to cocultures of 1D9-BBz T cells with MM.1s cells; **d**, cytolytic activity against luciferase-expressing (luc+) target cells, revealed as reductions in bioluminescence relative to target-only monocultures; and **e**, sustained cytolytic activity against fresh batches of CD3-CD138+ MM.1s cells added every 3 days, revealed by flow cytometric monitoring as delays in CD3-CD138+ cell expansion relative to untransduced T-cell cocultures. Each data point represents the mean of duplicate cultures using T cells from independent donors, with N indicating number of T-cell donors. Asterisks indicate significance per paired samples t test. **f, g, h, i.** Evaluation of 1D9-BBz T cells in preclinical mouse cancer models. **f,** Experimental workflow and treatment timeline. NSG mice bearing RPMI-8226 (P2RX5-FL/FL) tumors received a single intravenous (IV) infusion of 800,000 1D9-BBz T cells, untransduced controls, or cilta-cel (BCMA-directed CAR-T). Separately, NSG mice bearing T-ALL PDX models, either ETP-8 (P2RX5-FL/FL) or ALL-8 (P2RX5-FL/ΔC), were treated with a single IV infusion of 300,000 1D9-BBz T cells or untransduced controls. **g,** Longitudinal luc+ tumor growth assessed by bioluminescent imaging (top) and Kaplan–Meier overall survival curves (bottom). p-values were derived from the log-rank Mantel-Cox test. **h.** Representative images of tumor-bearing mice from the same cohorts. **i.** Flow cytometric enumeration of circulating human T-cell counts in the RPMI-8226 mouse model.

**Fig. 6.**
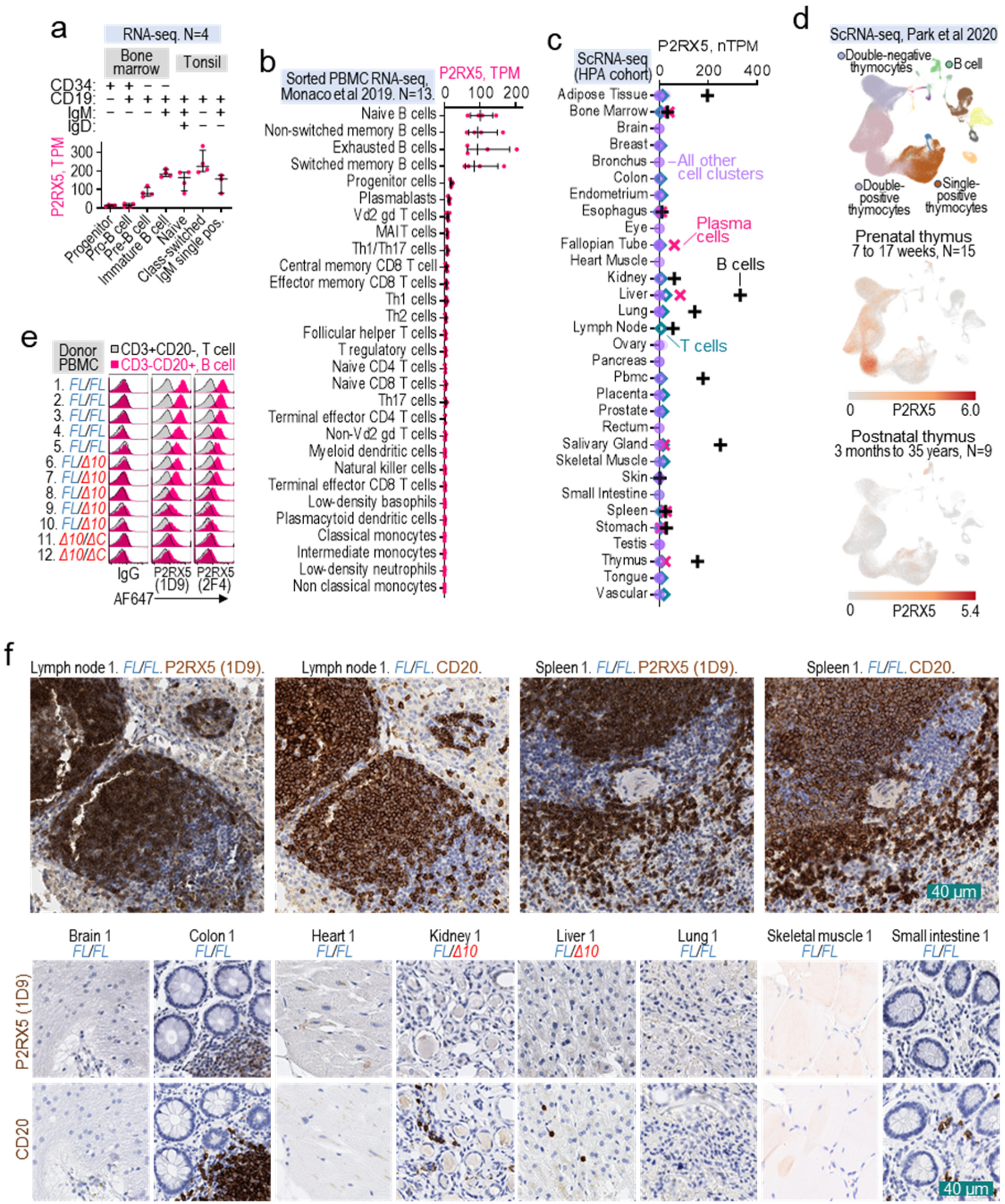
P2RX5 expression in postnatal tissues is B cell-restricted. **a-d.** *P2RX5* transcript levels as determined by bulk and scRNA-seq of human tissues: (**a**) bone marrow and tonsils [bulk]; (**b**) purified immune cell subsets from the peripheral blood [bulk]; **(c)** Human Protein Atlas (HPA) collection of post-mortem organs [scRNA-seq]; **(d)** pre- and postnatal thymuses [scRNA-seq]. In **a** and **b**, each data point represents an independent donor sample. In **c,** weighted means of *P2RX5* normalized transcripts per million (nTPM) among all cell clusters annotated as that specific cell type are shown. In **d,** UMAP projections are colored according to the expression level of the indicated gene. **e.** Live-cell flow cytometry of peripheral blood mononuclear cells (PBMCs) from healthy donors genotyped as *P2RX5 FL/FL*, *FL/Δ10* or *Δ10/ΔC*. Shown are staining intensities for IgG isotype control, 1D9, or 2F4 antibodies within populations gated as positive for either the CD3 T-cell or CD20 B-cell markers. **f,** Representative IHC images of normal tissue from donors expressing P2RX5 *FL/FL* or *FL/Δ10*. FFPE sections were stained for the expression of either P2RX5 (via 1D9 antibody) or CD20 using DAB substrate (brown), with hematoxylin nuclear counterstaining (blue).

We also parsed existing single-cell RNA-seq datasets corresponding to bone marrows from healthy donors ^32,33^ and from patients with large B-cell lymphoma ^34^, MM ^35^ or T-ALL ^36^ (**Extended Data Fig. 9a-d**), as well as all major human organs from Human Protein Atlas v24 ^28^ (**Fig. 6c**) and Tabula Sapiens ^37^ consortia (**Extended Data Fig. 9e, f**). We found that while high P2RX5 transcription is limited to CD20^+^ B cells in non-malignant postnatal tissues, this is not the case for prenatal organs. In fact, existing scRNA-seq data revealed that P2RX5 mRNA is comparably elevated in both thymocytes and B cells at prenatal stages, but thymic expression is rapidly extinguished months after birth, prior to thymus involution ^38^ (**Fig. 6d**).

To confirm the B-cell restricted expression of P2RX5 protein in adult lymphocytes, we performed live-cell flow cytometry of peripheral blood mononuclear cells (PBMCs) using 1D9 and 2F4 antibodies. We readily detected P2RX5 expression in CD20^+^ B cells, with the strongest staining of FL/FL, intermediate staining of FL/Δ10 and FL/ΔC, and no staining of Δ10/ΔC donor cells. Crucially, CD3^+^ T cells remained negative across all donors (**Fig. 6e**). Lastly, in IHC analysis of normal FFPE tissues, P2RX5 staining (via 1D9) was identical with that for CD20 in the lymph node, spleen and colon; whereas the brain, heart, kidney, liver, lung, and skeletal muscle were all negative (**Fig. 6f**; **Extended Data Fig. 10**).

## Discussion

The development of therapeutic targets is sometimes a non-linear, convoluted process. Over 20 years ago, P2RX5 (a.k.a. LRH-1) was identified as a minor histocompatibility antigen (MiHA) and shown to elicit an HLA-B7-restricted cytotoxic T cell (CTL) response in a chronic myelogenous leukemia patient infused with lymphocytes from a donor with a frameshift mutation in P*2RX5* ^1^. Of note, this allogeneic cytotoxic T-lymphocyte response induced a complete response without severe graft-versus-host disease, attesting to potential safety of P2RX5-directed immunotherapies. However, despite two follow-up studies ^39,40^, such therapies have not been developed clinically. In fact, the interest in *P2RX5* waned rapidly because of its perceived pseudogenization in humans ^2^. Here we show that not only its full-length, cell surface-bound variant (P2RX5-FL) is stably expressed in three-quarters of individuals of African ancestry, but it represents a potential target for CAR T cells (CART-X5) in a wide range of hematologic malignancies. This includes BCL and MM relapses which recurrently down-regulate targeted antigens [CD19 ^41–47^, CD22 ^48–56^, BCMA^57–61;62–66^, etc.] and immunotherapy-naive T-ALL.

P2RX5 expression in T-ALL is particularly unexpected given its absence in mature T cells. Although *ex vivo* activation was reported to upregulate the P2RX5 Δ10 variant in T cells ^23^, we detected only a modest increase for cell-surface P2RX5-FL that did not persist beyond a few days. Additionally, our survey of single-cell transcriptomic data found no significant P2RX5 induction across normal T-cell subsets *in vivo* and CAR-T cells during peak expansion in patients. However, unlike their post-natal counterparts, pre-natal thymocytes, especially CD4/CD8 double-positive (DP) cells, upregulate *P2RX5* mRNA to levels seen in prenatal B cells. What drives and later extinguishes this expression is currently unknown. One possible culprit is Notch1, a master regulator of T cell fate up to the DP stage ^67,68^. Notch1 and the associated oncofetal network could also be responsible for *P2RX5* upregulation in T-ALL, where Notch1 plays an important role in disease pathogenesis ^69,70^.

Irrespective of the underlying mechanism, selective activation of P2RX5 in T-ALL but not in normal T cells gives CART-X5 a key advantage over CD5- or CD7-directed CAR T cells, which may require extensive gene editing steps in both donor’s and recipient’s cells to prevent fratricide and post-infusion T-cell aplasia ^7,8^.Given its B cell-restricted expression in normal tissues, we expect P2RX5-targeted therapies to have similar safety profiles as therapies targeting B-cell markers, such as CD19 or CD20. We could not evaluate the on-target/off-tumor effects of CART-X5 in mouse models due to our antibodies lacking cross-reactivity with mouse P2rx5, combined with the distinct tissue expression pattern of mouse P2rx5 (including not seen in humans expression in brown adipocytes ^71^ and osteoclasts ^72^). We do, however, note that the aforementioned cytotoxic T-lymphocyte (CTL) response against P2RX5 induced complete remission without triggering severe graft-versus-host disease ^1^. While targeting a surface protein with CAR T cells is distinct from targeting its MHC-presented peptides with allogeneic CTLs, this clinical case study indirectly attests to the safety of P2RX5-directed therapies.

We also performed a high-throughput screening of 1D9 mAb against >6,000 human membrane proteins, which identified Nectin-4 as its sole off-target interactor. In our experimental system, 1D9 binding to Nectin-4 was weak and only apparent during ectopic Nectin-4 overexpression, and CART-X5 showed no signs of activation after co-culturing with keratinocytes and even Nectin-4 overexpressing carcinoma cell lines. Of note, existing IHC data indicate that Nectin-4 is absent across most postnatal tissues and is only expressed at low levels in keratinocytes, sweat glands, hair follicles, and transitional epithelium ^73,74^. Consistent with these findings, clinical trials of Nectin-4-directed antibody-drug conjugates (enfortumab vedotin ^74^) and CAR T cells ^75^ reported manageable on-target/off-tumor toxicities.

Finally, while P2RX5-directed therapies would be feasible only in selected populations of African ancestry, it is worth noting that the very same populations have benefited far less from modern forms of immunotherapy ^76^. In the U.S., only 1.3% of participants in clinical trials are of African ancestry even though they account for more than 25% of MM cases ^77,78^. Compared to non-African populations, African Americans also have two-fold higher incidence and mortality rate for MM ^79–81^ and worse survival for multiple forms of lymphoid malignancies, including CLL^82,83^, DLBCL^84^ and follicular and marginal zone lymphoma ^84^. It stands to reason that the advent of P2RX5-directed immunotherapies might help address some of these disparities.

## Methods

### RNA seq analyses

RNA-seq reads were first trimmed to remove adapters (BBTools v38.96) and were then aligned using STAR v2.7.9a to the hg38 reference genome while providing known gene isoforms through the GENCODE annotation V32. We used STAR flags “–quantMode GeneCounts” and “– alignSJoverhangMin 8” to quantify genes and ensure spliced reads had an overhang of at least eight bases. Junction spanning reads were obtained from the STAR “*_SJ.out.tab” result files, and each entry was normalized by dividing by the total number of junction-spanning reads and multiplying by a factor of 1 million to obtain the “junctions per million” or JPM. TPMs for all samples were calculated using TPMCalculator ^85^ version 0.0.3. Sample P2RX5 genotype were classified based on the ratio of [exon-10 to -11 JPM] / [exon-9 to -11 JPM] where values over 0.9 were classified as homozygous for exon-10 inclusion, in between 0.1 and 0.9 as heterozygous and below 0.1 as homozygous for exon-10 skipping. The rs3215407 alleles were determined via manual look-up of the alignments on IGV.

#### Bulk RNA-seq data usage

The Monaco PBMC (PRJNA418779) [Ref ^29^], Boultwood Bone Marrow (PRJNA473835) [Ref ^86^], bone marrow and tonsil subsets (PRJNA475684) [Ref ^26,27,53^], mouse and macaque tissue (GSE219045) [Ref ^87^], CCLE (PRJNA523380) [Ref ^88^], eBL Uganda 1 (PRJNA292327) and BL Uganda 2(PRJNA374464) [Ref ^15,16^], eBL West Kenya (phs001282.V2.p1) [Ref ^14^], MMRF CoMMpass (PRJNA248539) [Ref ^13^] datasets were from the BioProject or dbGaP databases of the National Center for Biotechnology Information (NCBI). For the CoMMpass dataset, samples were categorized as from donors of African ancestry based on prior principal component analysis (PCA) of 4,761 Ancestry Informative Markers (AIMs) SNPs ^89^.

### Single-cell transcriptomic data usage

ScRNA-seq-based normalized transcript counts per million (nTPM) for P2RX5 were downloaded from the Human Protein Atlas portal (HPA, version 24) ^37^. Within each dataset, we calculated the weighted mean nTPM counts for all clusters sharing a cell type, weighting by the number of cells per cluster. Clusters with mixed cell type or low-confidence cell type annotations were excluded.

### Cell culture

HEK293T and Jurkat cells were a kind gift from Dr. Peter Choi (Children’s Hospital of Philadelphia, Philadelphia, PA). DoHH2, Ramos, MM.1s, RPMI-8226 and MOLT-16 cells were obtained from the ATCC (American Type Culture Collection), while DND-41 and RPMI-8402 cells were from the Leibniz Institute DSMZ. Raji cells were authenticated with short tandem repeat profiling. All cell lines were routinely tested to be Mycoplasma free via EZ-PCR Mycoplasma Detection Kit (Sartorius). HEK293T cells were maintained in DMEM medium supplemented with 10% FBS, 2 mmol/L L-alanyl-L-glutamine dipeptide at 37°C and 5% CO2. All other cells were maintained in RPMI medium supplemented with 10% FBS, 2 mmol/L L-alanyl-L-glutamine dipeptide. To generate luciferase-zsGreen reporter cell lines, cells were transduced the pLC-Luc-IRES-zsGreen-P2A -Blast lentiviral vector and cultured in 10 ug/mL Blasticidin for 3 weeks.

### LCL generation

B95-8 cells (ATCC # CRL 1612; 13) were seeded at 3 x 10^5^ cells/ml in a 75cm^2^ tissue culture flask in complete RPMI 1640. After two days, cells were resuspended in fresh complete RPMI 1640 at 1 x 10^6^ cells/ml and stimulated with 20ng/ml tetradecanoyl phorbol acetate (TPA; Sigma-Aldrich) for 1 hour to induce virus production and then resuspended in complete RPMI 1640 for 96 hours before centrifugation at 600 g for 10 min at 4°C. Cell supernatant was then filtered through a 0.45-micron filter and stored at -80°C. For infection with EBV, 500µl of filtered supernatant containing EBV (B95.8) was added to 5X10^6^ PBMCs in 5 ml complete RPMI medium in a 25 cm^2^ tissue culture flask and placed upright in a CO_2_ incubator at 37°C for 2-4 hours. Subsequently, 10 ml additional medium and cyclosporin A (1µg/ml; Sigma-Aldrich, Inc., St. Louis, MO) was added. On days three and six, nine, and twelve, half of the medium was replaced with fresh complete RPMI with cyclosporin A. After two weeks, stable EBV+LCLs were formed, and anti-CD19 staining was used to confirm that the culture is 100% B cells by flow cytometry.

### DNA or RNA extraction

Total RNAs or genomic DNA were isolated from whole cells or FFPE sample scrolls using the appropriate Maxwell® RSC simplyRNA Cells, RSC RNA FFPE, RSC Cultured Cells DNA or RSC DNA FFPE Kits (Promega).

### Reverse transcription and PCR analysis

To generate cDNA, 500 ng of RNA (via NanoDrop™ 2000) were hybridized with a mixture of Oligod(T)_20_ and random hexamer primers and reverse transcribed with SuperScript™ IV VILO™ Master Mix (ThermosFisher Scientific). Traditional PCR was performed using the Q5® High-Fidelity 2X Master Mix (New England Biolabs). Primers include P2RX5 CDS (1332 bp amplicon on P2RX5FL cDNA; forward: ATGGGGCAGGCGGGCTGCAAGG; reverse: TCACGTGCTCCTGTGGGGCTC) and P2RX5 FLAG-KI (forward: GGTTACCAAGACGTCGACAC; reverse: CGACATCCCAGATCCGCTG). Quantitative PCR (qPCR) was performed using PowerSYBR Green PCR Master Mix (Life Technologies) on an Applied Biosystems Viia7 machine. According to the Minimum Information for Publication of Quantitative Real-Time PCR Experiments (MIQE) guidelines ^90^, all primers used had efficiencies exceeding 95% and amplified single PCR products from cDNA mixtures, which was then validated via Sanger sequencing. Reference gene primers used include RPL27 (123 bp amplicon on NM_000988; forward: ATCGCCAAGAGATCAAAGATAA; reverse: TCTGAAGACATCCTTATTGACG), GAPDH (227 bp amplicon on NM_002046; forward: GAAGGTGAAGGTCGGAGTC; reverse: GGAAGATGGTGATGGGATTTC) and ACTB (121 bp amplicon on NM_001101.5; forward: GTCATTCCAAATATGAGATGCGT; reverse: GCTATCACCTCCCCTGTGTG) ^91,92^. P2RX5 primers: exon-10 skipping (Δ10) (93 bp amplicon; forward: TGATGGTGAACGGCAAGGGT; reverse: CCTCGTACTTCTTGTCACGG), exon-10 inclusion (∇10) (135 bp amplicon; forward: AGTTCAGCATCATTCCCACCA; reverse: CCTCGTACTTCTTGTCACGG) and exon 9 (pan) (55 bp amplicon; forward: ATATTACCGAGACGCAGCCG; reverse: CCCGTAGGCTTTCATCAGGG). Expression levels relative to control were calculated using the 2-ΔCT and 2-ΔΔCT methods. The data were normalized to the CT or RPL27 or when indicated, the mean CT of GAPDH, ACTB and RPL27. DNA contamination was routinely assessed via qPCR of negative control reverse transcription reactions which lacked reverse transcriptase (cycle thresholds above 35).

### Plasmid constructs

The *P2RX5* minigene sequence (exons 9-11) containing truncated introns and either the rs891746(A or C) alleles was fused upstream of a NanoLuc ORF and cloned into the MXS_CMV::PuroR-bGHpA plasmid, replacing the PuroR ORF. MXS_CMV::PuroR-bGHpA was a gift from Pierre Neveu (RRID: Addgene_62439) ^93^. The pCDH-EF1 vector was a gift from Kazuhiro Oka (RRID: Addgene_72266). The pLC-Luc-IRES-zsGreen-P2A -Blast lentiviral vector was constructed by replacing the CMV promoter sequence (immediately upstream of the zsGreen ORF) in pLC-ZsGreen-P2A-Blast with the EF1a promoter, Luciferase ORF and IRES sequence of pHIV-Luc-ZsGreen. The pLC-ZsGreen-P2A-Blast plasmid was a gift from Eva Gottwein (RRID: Addgene_123322) (PMID: 29954751) while pHIV-Luc-ZsGreen was a gift from Bryan Welm (RRID: Addgene_39196).

### Gene knockouts

Alt-R™ CRISPR-Cas9 sgRNAs (Integrated DNA Technologies) were complexed with Alt-R™ S.p. Cas9 Nuclease V3 (Integrated DNA Technologies). The gRNA sequences targeting CD19 (CTAGGTCCGAAACATTCCAC), TNFRSF17 (TAACGCTGACATGTTAGAGG), P2RX5 (TATATCCTGGAAGTCGCTCC and AAGGACAGCGACTGCCACGC) and CD38 [described previously ^94^] were used. The resulting ribonucleoprotein complexes were transfected into the target cells using the Neon Electroporation System (Invitrogen).

### P2RX5 variant determination

For donor samples and cell lines, both genomic DNA and cDNA was genotyped using predesigned Taqman probes (Thermo Fisher Scientific. C_8728453_10 for rs891746.). RT-qPCR was also performed to determine if pan P2RX5 mRNA was upregulated, and if exon-10 was included or skipped. For cell lines, genomic DNA was also genotyped using PCR amplification followed by Sanger or Illumina® Short-read NGS sequencing for rs3215407 (251 bp amplicon; forward: GCCCTCTTTCAGGTAACGCT; reverse: GGAGTGGACAGACACCCTTG) and rs891746 (231 bp amplicon; forward:TGGGGGCCCCGCTGTCCAAC; reverse:TCTTTCTGGGGGCACTGGGACAC).

### Splicing minigene assay

The minigene plasmids were electroporated into Raji and OCI-ly8 cells using the Neon System (Invitrogen). Two days later, minigene transcription was analyzed via RT-qPCR and Sanger sequencing using the “minigene primers” [(495 bp amplicon; forward: ATATTACCGAGACGCAGCCG; reverse: GCCAGTCCCCAACGAAATCT)].

### Antibody generation

Anti-P2RX5 monoclonal antibodies were generated at the Fred Hutchinson Antibody Technology Core. Briefly, male and female 20-week-old mice were immunized with NIH/3T3 cells that stably overexpressed P2RX5FL-FLAG. Following a 12+ week boosting protocol, splenocytes were isolated from high-titer-yielding mice and electrofused with FOX-NY myeloma cells. From the resulting hybridoma clones, two unique monoclonal antibody sequences, 1D9 and 2F4, were reactive to cell-surface P2RX5-FL in flow cytometry assays. 1D9 and 2F4 were produced in serum free media and purified using protein A and protein G columns on AKTA pure HPLC.

### Western blotting

Total cell lysates were prepared in Laemmli Sample Buffer (Bio-Rad) with 50 mM DTT, resolved on a 4–15% precast polyacrylamide gel (Mini-PROTEAN® TGX™, Bio-Rad) in Tris/glycine/SDS running buffer (Bio-Rad), and then transferred onto a PVDF membrane (Millipore Sigma, Immobilon-P, 0.45 µm) via the Mini Trans-Blot® Cell system (Bio-Rad). These membranes were then probed with antibodies against P2RX5 (1D9 clone), FLAG (L5 clone, Biolegend), CD79B (D7V2F clone, Cell Signaling Technology), ACTB (8H10D10 clone, Cell Signaling Technology), GAPDH (14C10 clone, Cell Signaling Technology); followed by horseradish peroxidase–linked secondary antibodies (Cell Signaling Technology). Western blot chemiluminescent signals were captured with a G:Box Chemi XX6 Gel Doc (Syngene).

### Cycloheximide chase assay

Cell lines were transduced with lentiviral particles ORFs for the indicated constructs or an empty vector control followed by 2-week selection with 10 μg/mL Puromycin (Invitrogen). Cultured Raji cells were treated with 60 µg/mL of cyclohexmide (CHX) (Sigma-Aldrich, C1988-1G).

### Protein deglycosylation

Raji and MM.1s cell lysates (in RIPA buffer) were deglycosylated with the Protein Deglycosylation Mix II (P6044, New England Biolabs), under denaturing reaction conditions, according to the manufacturer’s protocol. The Protein Deglycosylation Mix II was replaced with water for the negative control reaction.

### Human samples

Archival FFPE samples were obtained from the Penn Tumor Tissue/Biospecimen Bank (TTAB) under the Penn Institutional Review Board (IRB) approved protocol 856585. Cryopreserved bone marrow aspirates from Black or African American patients with T-ALL were obtained from the CHOP Center for Childhood Cancer Research Biobank and Repository under the CHOP IRB-approved protocol #10-007767. Bone marrow aspirates from Black or African American patients with MM were collected under the approved IRB protocol 842940 through the PCD group at the Hospital of the University of Pennsylvania.

### Immunohistochemistry (IHC)

Formalin fixed, paraffin embedded (FFPE) section staining was performed on a Bond Max automated staining system (Leica Biosystems). Antigen retrieval was performed for 20 mins with the E2 (CD20 and CD138 staining) or E1 (P2RX5 staining) retrieval solution (Leica Biosystems). For CD20 (clone L26, Dako) and CD138 (clone MI15, Dako) staining, we used the Bond Refine polymer staining kit (Leica Biosystems) with 1 hour and 15 mins room temperature primary antibody incubation, respectively. For P2RX5 staining, we used the ImmPRESS Excel Amplified Polymer Staining Kit, Anti-Mouse IgG, Peroxidase (MP-7602-15; Vector Laboratories) with 1 hour room temperature primary antibody (clone 1D9) incubation. Slides were rinsed, dehydrated through a series of ascending concentrations of ethanol and xylene, and then cover slipped. Stained slides were then digitally scanned at 20x magnification on an Aperio CS-O slide scanner (Leica Biosystems). Representative pictures were taken on scanned IHC slides with Aperio ImageScope program v12.2.2.5015.

### Live-cell flow cytometry

Live cells (cell lines, patient bone marrow aspirates or mouse peripheral blood) were incubated in blocking buffer (PBS with 2% FBS, 1 mM EDTA and Human TruStain FcX™ (Biolegend)) for 20 min prior to 30 min staining with one or a combination 1D9, 2F4 or Biolegend antibodies against: FLAG, L5 clone; CCR7, G043H7 clone; CD19, HiB19 clone; CD20, 2H7 clone; CD22, HIB22 clone; CD38, HIT2 clone; CD45(human specific), H130 clone; CD45RA, HI100 clone; CD79B, CB3-1 clone; BCMA, 19F2 clone; CD3, UCHT1 clone; CD7, 4H9 clone; and CD138, DL-101 clone. When unlabeled 1D9 or 2F4 primary antibodies were used, the blocking buffer was supplemented with 10% (v/v) goat serum (Fisher Scientific, 16-210-064) and an additional 30 min secondary antibody staining with Goat anti-Mouse IgG (H+L) Cross-Adsorbed Secondary Antibody, Alexa Fluor™ 647 (Invitrogen, A-21235), was performed. For competition assays, the blocking buffer was supplemented with 80 ug/mL of unlabeled 1D9 or 2F4 antibodies. All incubations were at 4°C. Acquisition was done on a BD Accuri™ C6 Cytometer (BD Biosciences). Manual gating was performed on FlowJo Software v.10. Briefly, a FSC vs SSC density plot was first used to exclude cell debris followed by doublet exclusion on a FSC-H vs FSC-A plot. Subsequently, propidium Iodide (PI) positive dead cells were excluded from analysis before marker staining was plotted on a histogram (for a single marker) or a density plot (for 2 markers). Boundaries for “positive” and “negative” populations were defined based on the unstained control samples (for single marker stains) or the fluorescence minus one (FMO) controls (two marker stains). Median fluorescence intensity (MFI).

### Multiparameter flow cytometry

Cells were resuspended in FACS staining buffer (phosphate buffered saline (PBS) +2% FBS) using a combination of the following antibodies against: CD14, clone M5E2; CD4, clone OKT4; CD8ECD, clone B9.11; CD3, clone OKT3; CD34, clone 561; CD45, clone 2D1; CD5, clone BL1a; CD7 CD7-6B7, P2RX5, clones 1D9 and 2F4. Cell viability was established using Zombie UV Dye (Biolegend), and data were acquired on Cytek Aurora (Cytek Biosciences). Analysis was performed using FlowJo 10.8.0 software (FlowJo, LLC).

### Manufacturing of CAR T-cells

Healthy donor PBMCs were obtained from the Human Immunology Core at the University of Pennsylvania or from STEMCELL Technologies. T-cells were isolated from frozen donor PBMCs using the EasySep™ Human T-Cell Enrichment Kit (STEMCELL Technologies). Isolated T-cells were cultured in ImmunoCult™-XF T Cell Expansion Medium (STEMCELL Technologies) supplemented with 100 IU/ml of recombinant human IL-2 (Preprotech), and activated with ImmunoCult™ Human CD3/CD28 T Cell Activator (STEMCELL Technologies). Lentiviral transduction was performed the next day. The number viable cells in culture was assessed every 2 to 3 days using a Countess 3 Automated Cell Counter with tryphan blue staining and the number of population doublings was calculated. At day-7 post CD3/CD28 activation, T-cells were cryopreserved. The % of CAR positive T-cells was monitored via flow cytometry analysis upon staining with the MonoRab™ Rabbit Anti-scFv Cocktail [iFluor 647] (Genscript).

For the 1D9-BBz CAR T cells and ciltacabtagene autoleucel used in the mouse xenografts models, the CAR T cells were generated as previously described ^95^. Briefly, CD4+ and CD8+ cells were combined at a 1:1 ratio and activated using anti-human CD3/CD28 Dynabeads (Invitrogen, #40203D) at a 3:1 ratio. 24 hours (day 1) post activation, T cells were infected with lentiviral vectors containing the CAR transgenes (multiplicity of infection ∼1.5 viral particles / T cell). Magnetic beads were removed from T cells on day 5.

### IFNG secretion assay

Target cells were co-cultured with expanded healthy donor T cells at day 12 post CD3/CD28 activation, at 1D9-BBz effector CAR T cell to target cell (E:T) ratio of 1:2 for a day before media IFNG levels were measured using Lumit® IFN-γ (Human) Immunoassay (Promega) and a FLUOstar Microplate Reader (BMG Labtech). IFNG levels were normalized to the MM.1s with 1D9-BBz CAR T cell coculture.

### Cell-to-cell avidity assays

Avidity was measured using the Lumicks Z-Movi assay. Z-Movi-compatible acoustofluidic chips were coated with poly-L-lysine for 3 h prior to attaching a monolayer of RPMI-8226 cells. CellTrace far-red labeled CAR T cells were allowed to incubate on the monolayer of RPMI-8226 for 10 min, and then a ramping acoustic force was applied. Cell-detachment was analyzed using ImageJ and R. Avidity experiments were conducted according to the manufacturer’s (LUMICKS™) instructions.

### Luciferase reporter-based in vitro killing assays

Luciferase-expressing (luc+) target cells were co-cultured with expanded healthy donor T cells at 9-days post CD3/CD28 activation at the indicated effector CAR T cell or T cell to target cell (E:T) ratios for 1 or 4 days before luciferase activity was monitored with the ONE-Glo™ EX Luciferase Assay System (Promega) and a FLUOstar Microplate Reader (BMG Labtech), as a proxy for cell viability. The luminescence signal was normalized to target cell alone monocultures.

### Repeat tumor cell challenge assay

Expanded T cells at day 12 post CD3/CD28 activation were co-culture with target cancer cells. Fresh batches of MM.1s cells were added every 3 days and viable, propidium iodide-negative CD3-CD138+ cell counts were determined via live-cell flow cytometry. Cell counts were normalized with CountBright™ Absolute Counting Beads (Thermo Fisher Scientific) and reported as the number of population doublings.

### Fratricidal activity assay

Unlabeled effector T cells (1D9-BBz or untransduced controls at day 9 post CD3/CD28 activation) were cocultured with one of the following labeled target cells (Thermo Fisher CellTrace™ Far Red Cell Proliferation Kit): 1. T cells from the same FL/FL donor activated 3 days prior; 2. the same batch of T cells except non-activated; and 3. MM.1s cells. The propidium iodide negative, CellTrace-positive target cell populations were counted daily, over 6 days via flow cytometry. Cell counts were normalized with CountBright™ Absolute Counting Beads (Thermo Fisher Scientific) and reported as the number of population doublings.

### Animal models

Mice studies were carried out in accordance with Children’s Hospital of Philadelphia and University of Pennsylvania Institutional Animal Care and Use Committee (IACUC) guidelines. To compare the 1D9-BBz CAR T cells with ciltacabtagene autoleucel, 6- to 8-week-old NSG mice were injected intraosseously with RPMI-8226-luciferase cells as described earlier ^95^. A week later, mice were injected intravenously with a total of 0.8×10^6^ CAR T cells. For the T-ALL PDX models, 4 month-old NSG mice were injected intravenously with ETP-8- or ALL-8-luciferase-GFP cells as described earlier ^7^). After 10 days, mice were injected intravenously with a total of 0.3×10^6^ CAR T cells. Whole animal bioluminescent imaging was performed with the IVIS Lumina S5 (PerkinElmer) and the images were analyzed using Living Image, v.4.1, software (PerkinElmer). The total bioluminescent signal flux for each mouse (flank for RPMI-8226; ventral for ETP-8 and ALL-8) was expressed as average radiance (photons per second per square centimeter per steradian). Survival was recorded and analyzed at the end of the study. For the RPMI-8226 model, peripheral blood was collected from all animals and the absolute numbers of live human CD45+ murine CD45-CD3+ T cells were determined by flow cytometry.

### Statistics

Statistical differences between two groups were evaluated using the paired t test. Kaplan-Meier survival curves were evaluated using Logrank (Mantel-Cox) test. P values were categorized as: > 0.05 (ns), ≤ 0.05(*), ≤ 0.01(**) and ≤ 0.001(***). All statistics were performed in Prism software (v.11.1, GraphPad).

