## Supplemental Figures 1-10 for "Ancestry-restricted expression of P2RX5 defines a CAR T-cell target across diverse blood cancers"

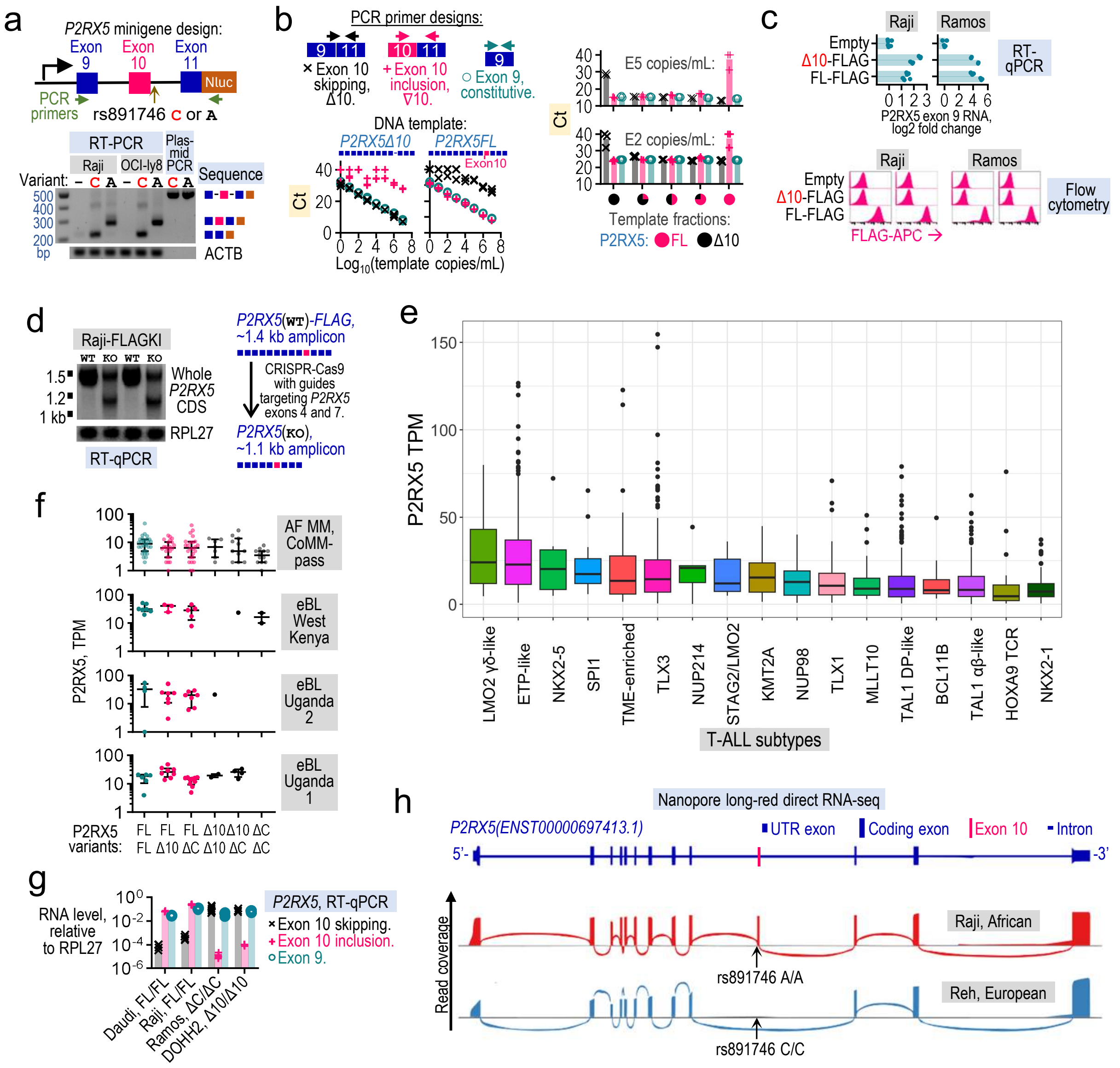


**Fig. 1 | Full-length P2RX5 (P2RX5 FL) mRNA is expressed across multiple lymphoid malignancies. a.** Splicing minigene assay. Top: Design of minigene expression vectors containing the sequence of *P2RX5* from exons 9 to 11, with truncated introns and either the rs891746 A or C alleles. PCR primer bindings sites are shown in green. Bottom: RT-PCR analysis detecting a ~300 bp mRNA amplicon (which includes exon 10) in Raji and OCI-ly8 cells after transfection with the rs891746 A allele-containing minigene, while cells transfected with the rs891746 C allele-containing minigene produce a ~220 bp mRNA amplicon (which skips exon 10). Control PCRs used the minigene expression plasmids as templates. Amplicons were separated by agarose gel electrophoresis and stained with EtBr. On the right of the gel image, are representations of the Sanger sequencing results for the indicated amplicons. **b.** Primer pair bindings sites for the detection of P2RX5 exon 10 skipping or inclusion, and exon 9 which is a constitutive exon found in all P2RX5 isoforms (top left). These primers accurately quantified single (bottom left) and mixed (right) dsDNA template standards in real-time quantitative PCR assays (qPCR). Ct: Cycle threshold. **c.** Analysis of Raji and Ramos cells after transduction to overexpress P2RX5 FL or Δ10 with extracellular FLAG tags. Top: RT-qPCR detecting the overexpression of both FL and Δ10, as an upregulation in the exon 9 sequence found in both FL and Δ10 transcripts. Each bar represents a replicate cell sample. Bottom: The FL variant, but not Δ10, was detected on the cell-surface as an increased in FLAG staining, during live-cell flow cytometry. Data from two replicate experiments are shown. **d.** RT-PCR analysis of Raji FLAG tag knock-in (FLAG-KI) cells showing a reduction in P2RX5 transcripts containing the full-length (~1.4 kb) coding sequence after a second modification to knockout (KO) *P2RX5*. PCR amplicons were separated by agarose gel electrophoresis and stained with EtBr. **e.** RNA-seq analysis showing *P2RX5* abundance (Transcripts Per Million, TPM), across T-ALL subtypes in the COG AALL0434 clinical trial cohort (N = 1309). **f.** RNA-seq analysis showing elevated *P2RX5* abundance (TPM), in MM patients of African ancestry (MMRF CoMMpass cohort) and endemic Burkitts lymphoma (eBL) patients from Africa. Samples were stratified according to the P2RX5 variants expressed. **g.** RT-qPCR analysis detecting *P2RX5* mRNA with exon 10 included for Daudi and Raji cells, and with exon 10 skipped for Ramos and DoHH2 cells; while the constitutive exon 9 was detected across all cells. **h.** Nanopore long-read direct RNA sequencing showing exon 10 inclusion by the P2RX5 mRNA transcripts detected in rs891746 A homozygous Raji cells, and exon 10 skipping in rs891746 C homozygous Reh cells. Top: MANE annotations of *P2RX5* exon-intron structure, in blue. Bottom: Sashimi plots showing read densities across all *P2RX5* exons. Arcs that connect exons represent splice junctions. Sanger sequencing results for the rs891746 alleles are indicated.

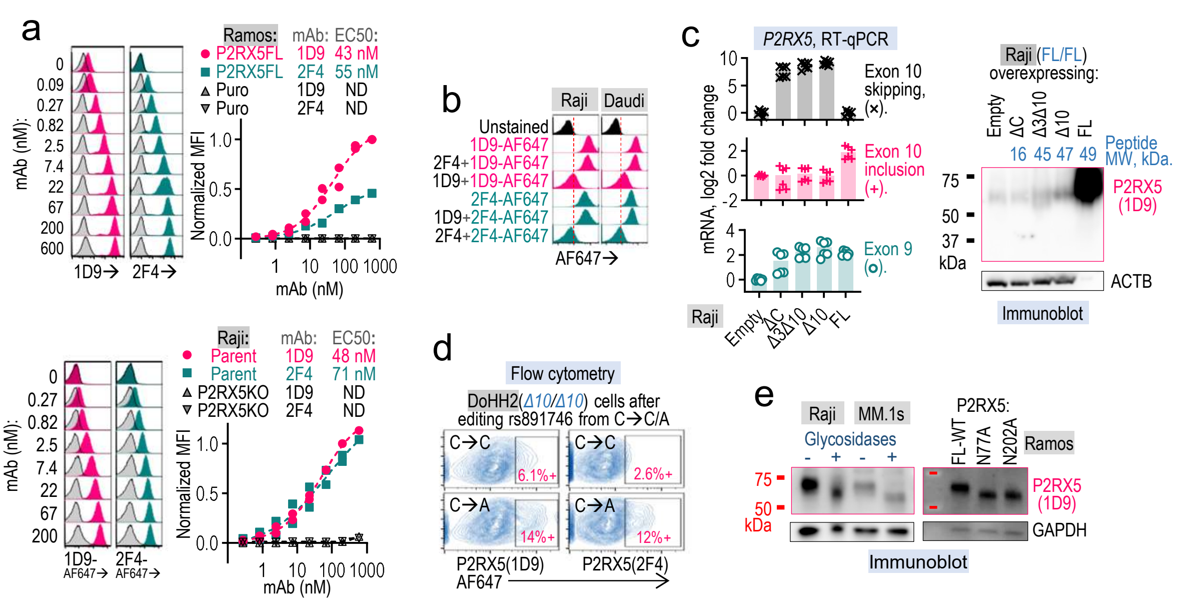


**Fig. 2 | P2RX5 FL protein expression in cell lines. a, b.** P2RX5-FL expression in cell lines, as detected via live-cell flow cytometry. **a,** Antibody titration assays: Ramos cells overexpressing empty vectors (Puro) or P2RX5FL, as well as Raji parent and P2RX5 knockout (P2RX5KO) derivates, were stained with unlabeled 1D9 or 2F4 antibodies at the indicated concentrations prior to indirect staining with fixed concentrations of AF647-labelled secondary antibody. **b,** Antibody competition assays: Raji and Daudi cells were pre-incubated with unlabeled 1D9, 2F4 or no blocking antibody, before staining with Alexa Fluor 647 (AF647) labeled 1D9 or 2F4. **c.** P2RX5 expression in Raji cells upon transduction to express P2RX5 ΔC, Δ3Δ10, Δ10 or FL variants, or an empty vector control. Left: RT-qPCR showing an increase in exon 9 mRNA found across all *P2RX5* variants, while only transduction with the FL variant led to an increase in exon 10. Values normalized to empty vector control cells. Right: Immunoblots using 1D9 antibody detect a greater abundance of FL protein, relative to the ΔC, Δ3Δ10 and Δ10 variants. The FL-overexpression lane was loaded with 90% less total cellular protein compared to other lanes to avoid signal saturation. Shown in blue are the calculated molecular weights (MW) of each overexpressed variant based on amino acid sequence alone. **d.** Live-cell flow cytometry. DoHH2 cells were subject to CRISPR-Cas9-mediated genome editing at rs891746 from C-to-C (C→C) or from C-to-A (C→A). This led to an increase in cell-surface P2RX5, as detected via the 1D9 and 2F4 antibodies. **e.** Representative immunoblots with the 1D9 antibody showing reduced P2RX5 protein sizes (in kDa) after Raji and MM.1s cell lysates were treated with (+) glycosidases, compared to untreated (-) controls (left); and in Ramos cells overexpression mutant variants of P2RX5FL with N77A or N202A substitutions, compared to cells expressing the wildtype (WT) protein.

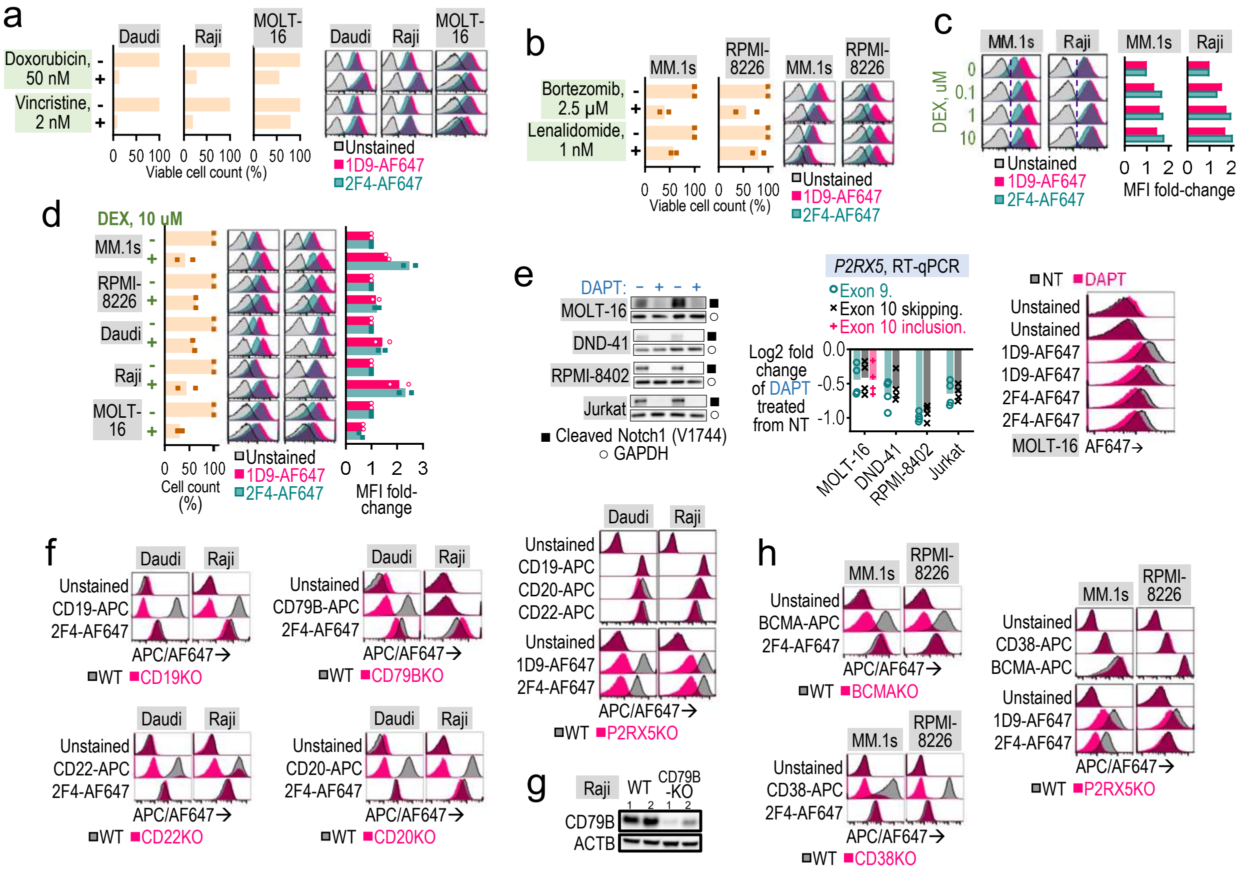


**Fig. 3 | P2RX5 FL expression in cell lines persists after chemotherapy drug exposure and immunotherapy target knockout. a-e.** Live-cell flow cytometric analysis showing positive P2RX5 expression (via 1D9 and 2F4 antibody staining) in cell lines after treatment (+) for 3-days with the indicated compounds. The compound concentrations reduced the number of viable, trypan blue-negative cells (as assessed via automated cell counter), relative to untreated (-) controls. **e,** Left: Immunoblots showing Notch1 cleavage at Valine1744 being inhibited by DAPT treatment. Middle: RT-qPCR showing the log2 fold change of P2RX5 transcript downregulation in DAPT treated cells, relative to untreated (-) cells. **f, h.** Live-cell flow cytometric analysis showing that the indicated cell lines do not lose P2RX5 positivity after CD19, CD20, CD22, CD79B, CD38 or BCMA knockout, and vice versa. **g.** Representative immunoblot of CD79B expression since CD79B expression on Raji cells is predominantly intracellular with little to no cell-surface protein.

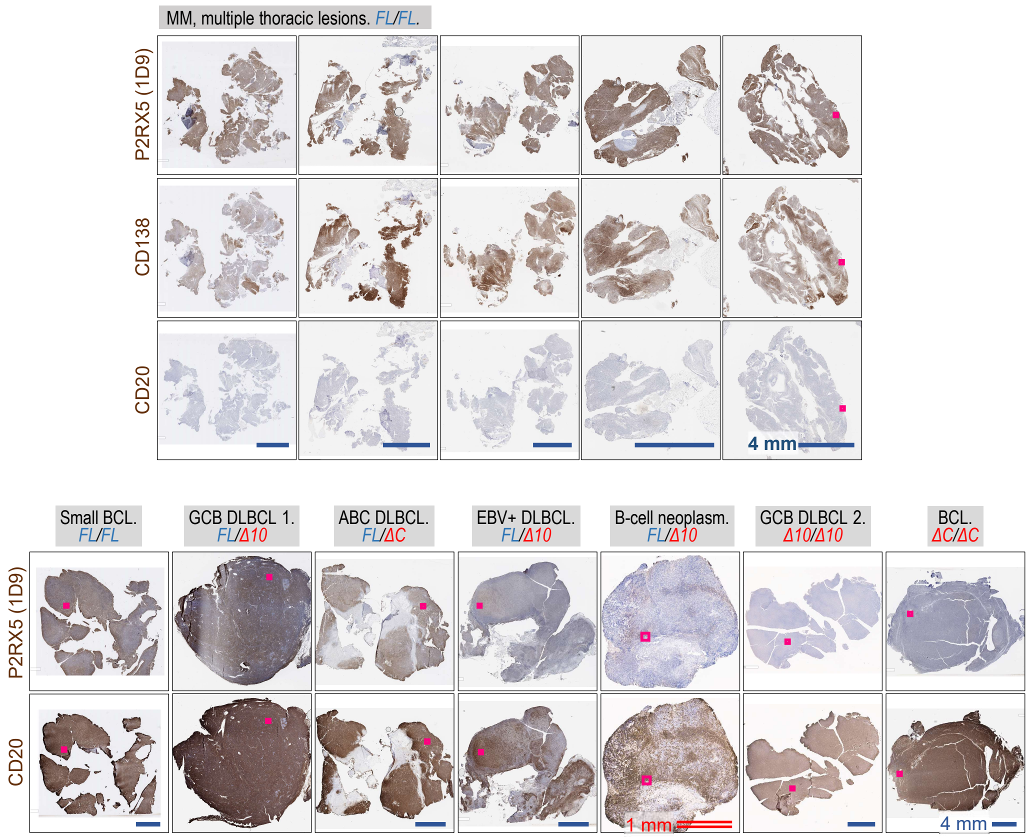


**Fig. 4 | Only BCL and MM tumor lesions with the P2RX5 FL haplotype stain positive for P2RX5 protein.** IHC images of whole FFPE tumor lesions, stained as indicated for P2RX5 (via 1D9 antibody), CD138 or CD20 using DAB substrate (brown), and with hematoxylin nuclear counterstaining (blue). Images of zoomed-in areas, as marked by red squares, are shown in Main Fig. 4a.

Main Fig. 4c example: **MM**, ΔC/ΔC. IgG-AF647 stained control.

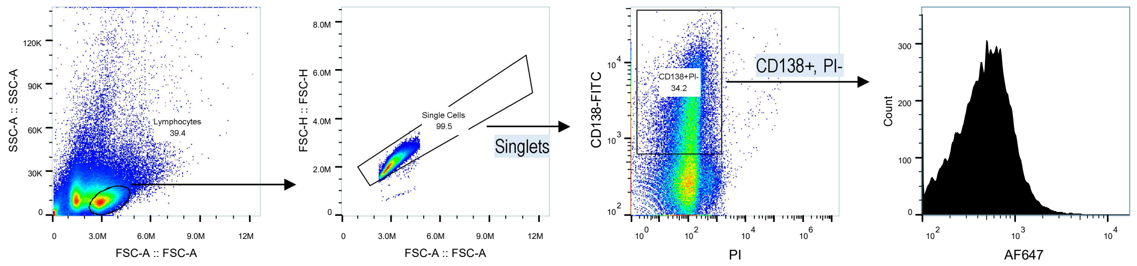


Main Fig. 4c, bottom panel example: **T-ALL**, FL/ΔC. Flourescence minus one (FMO) control.

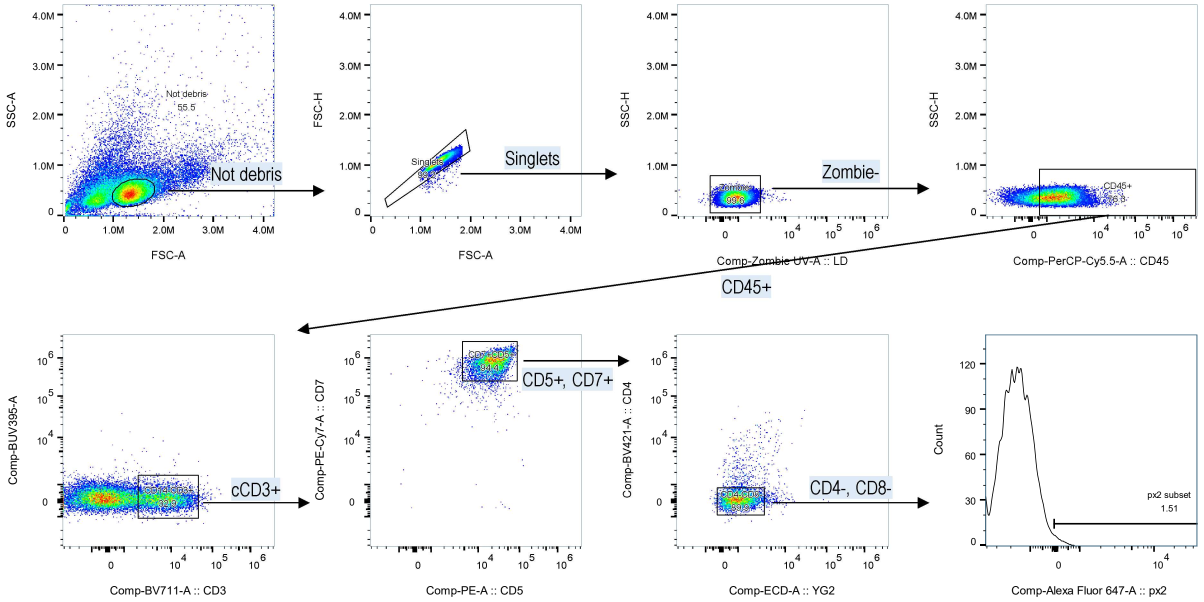


Main Fig. 4d, left panel example: **ETP8**, *FL/FL*. IgG-AF647 stained control.

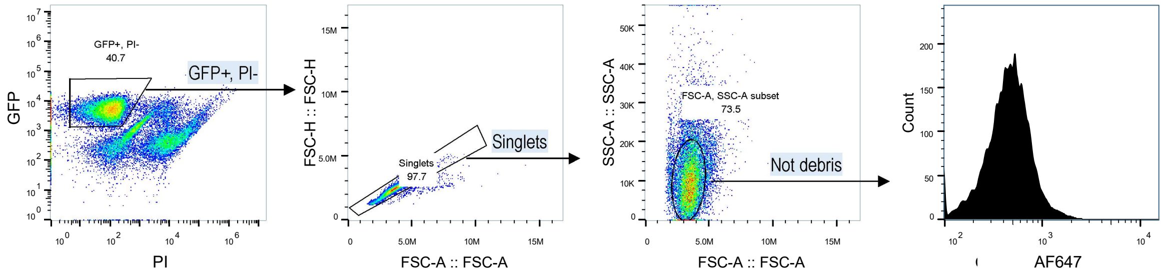


Main Fig. 4d, right panel example: **TH33**, Δ10/ΔC. IgG-AF647 stained control.

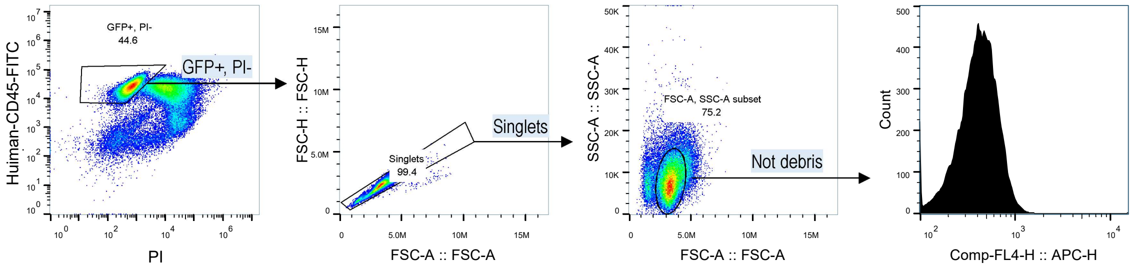


**Fig. 5 | Flow cytometry example gates for Main Fig. 4c, d.** GFP or human CD45 expression distinguishes xenografted human T-ALL cells from murine cells.

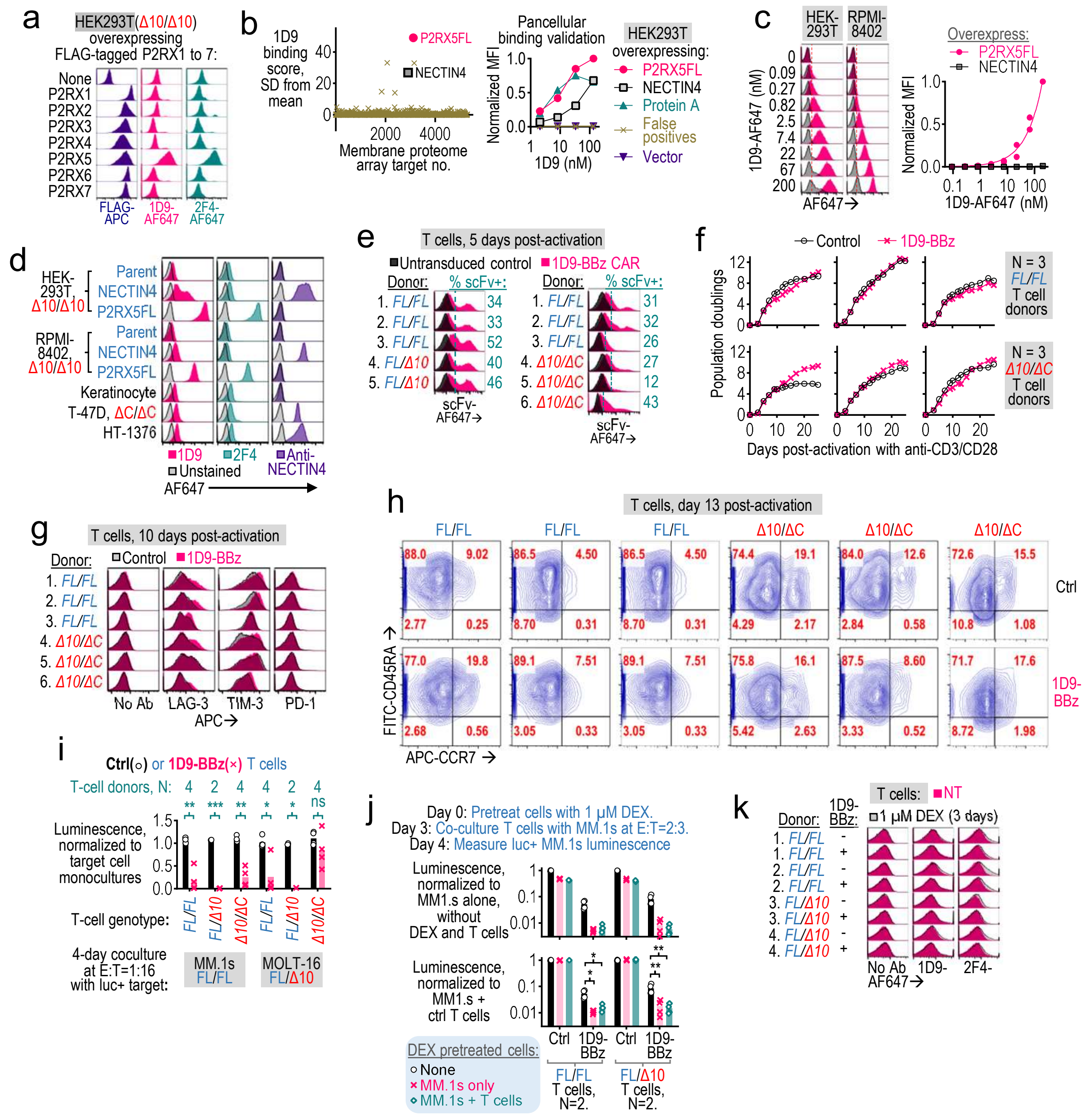

**Fig. 6 | 1D9-BBz CAR-T cells display enhanced cytolytic activity against P2RX5 FL-expressing tumors. a.** The 1D9 and 2F4 antibodies stain HEK293T cells overexpressing P2RX5FL-FLAG but not other FLAG-tagged P2RX family proteins, as assessed via live-cell flow cytometry. All P2RX proteins reached comparable cell-surface expression levels, based on FLAG tag staining. **b.** 1D9 tested against >6,000 human membrane proteins in unfixed but permeabilized HEK293T cells, using the Integral Molecular Membrane Proteome Array. NECTIN4 was the only off-target interaction detected. **c, d.** Confirmatory live-cell flow cytometry analysis showing 1D9 staining that is strong for HEK293T-P2RX5FL and RPMI-8226-P2RX5FL cells which stably overexpress P2RX5 FL, weak for HEK293T-NECTIN4 and RPMI-8226-NECTIN4 cells which stably overexpress Nectin-4, and comparable to 2F4 background staining for carcinoma cell lines with endogenous Nectin-4 overexpression (T-47D and HT-1376 cells). **e.** Representative flow cytometry histograms showing transduction efficiencies 1D9-BBz CAR into donor T cells (N=8). Transduced cells expressed cells-surface 1D9 scFv. **f.** Expansion of T cells during ex vivo activation with soluble anti-CD3/CD28, as assessed via automated counting of viable (tryphan blue negative) cells every 2-3 days. 1D9-BBz CAR transduced T cells expanded at comparable rates to the untransduced controls. **g, h.** Live-cell flow cytometry analysis of the 1D9-BBz CAR T-cell products. For **g**, LAG-3, TIM-3 and PD-1 T-cell exhaustions marker expression were not increased in 1D9-BB T cells relative to untransduced controls. For **h**, the ratio of CD45RA+CCR7+ naïve T cells (TN), CD45RA-CCR7+ central memory T cells (TCM) and CD45RA-CCR7- effector memory T cells (TEM) populations. **i, j**. Compared to untransduced controls, coculture with 1D9-BBz T cells led reductions in the number of luciferase-expressing (luc+) MM.1s or MOLT-16 target cells, which was detected as a reduction in bioluminescence. For **i**, after 4-day coculture at 1:16 effector-to-target (E:T) ratio. For **j**, MM.1s or effector T cells or both were pretreated for 3-days with 1 µM dexamethasone (DEX), before DEX wash-out and coculture. **k.** Cell-surface P2RX5 levels, as assessed via live cells flow cytometry with 1D9 and 2F4, in DEX treated (+) T cells relative to untreated (-) controls.

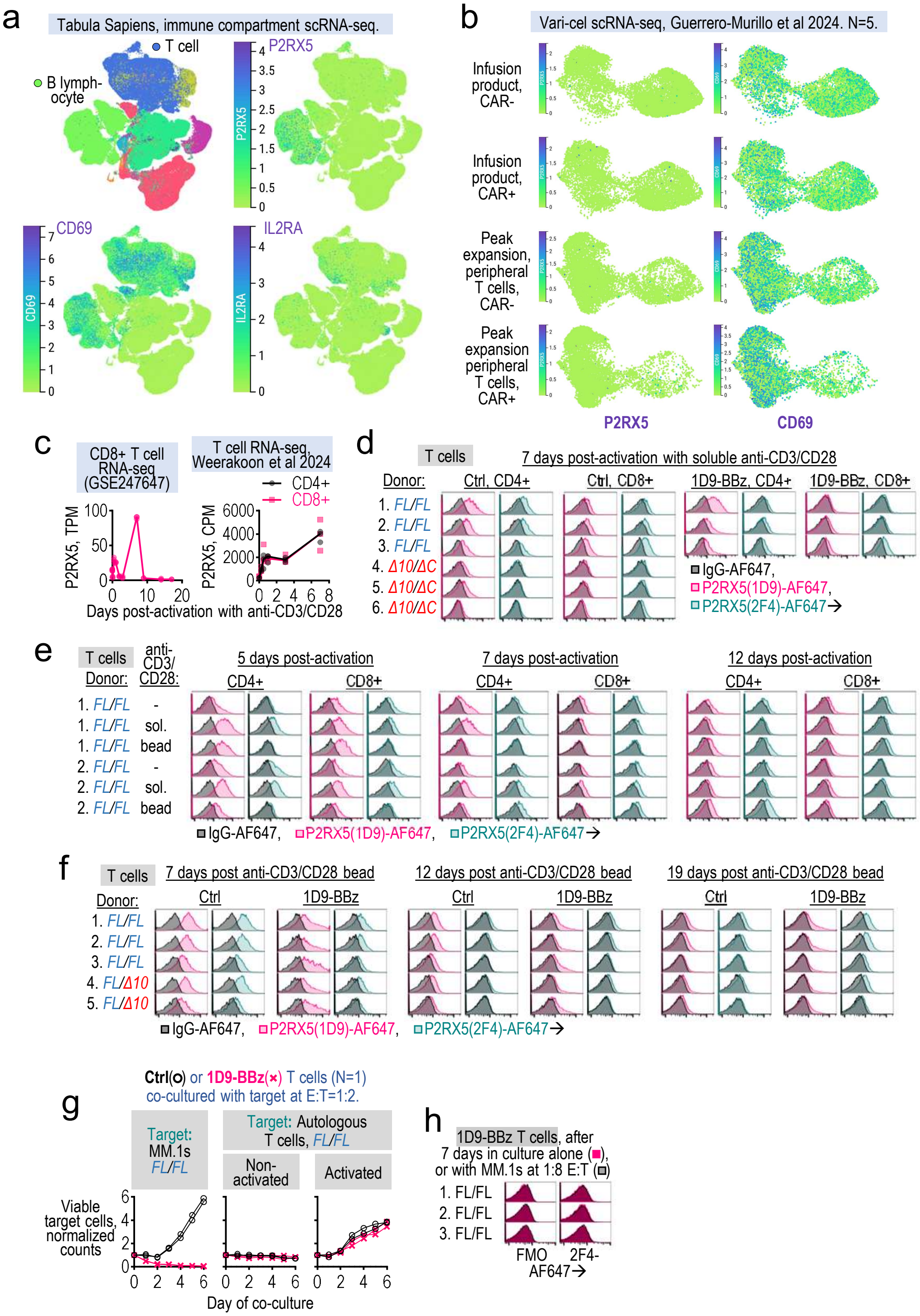


**Fig. 7 | T cells lack of sustained P2RX5 expression and are resistant 1D9-BBz CAR lysis. a, b.** Single cell transcriptomics data of P2RX5 expression in the immune compartment of 28 human organs from the Tabula Sapiens v2 portal (**a**), and of CD19-directed CAR-T cells (varnimcabtagene autoleucel or var-cel), in the infusion product (IP) and in peripheral blood at the time of peak CAR-T cell expansion following infusion (**b**). T cells show minimal P2RX5 expression, while upregulating CD69 and IL2Ra T-cell activation markers. UMAP projections are colored according to the expression level of the indicated gene. **c.** Longitudinal RNA-seq analysis showing transient induction of P2RX5 transcription in CD4+ and CD8+ T cells, after ex vivo activation with anti-CD3/CD28 beads (reanalyzed from GSE247647 and Weerakoon et al. 2024). **d-f.** Live-cell flow cytometry analysis showing transient inductions of cell-surface P2RX5 levels in cultured T cells following activation with soluble or bead-bound anti-CD3/CD28 antibodies. P2RX5 expression (detected via 1D9 and 2F4 antibodies), relative to matched IgG isotype controls. T cells from 8 independent donors representing distinct P2RX5 genotypes: FL/FL, FL/Δ10 and Δ10/ΔC. **g.** Unlabeled effector T cells (1D9-BBz or untransduced controls) were cocultured with one of the following CellTrace-labeled target cells: 1. T cells from the same *FL/FL* donor activated 3 days prior; 2. the same batch of T cells except non-activated; and 3. MM.1s cells. The expansion of the labeled target cell populations were monitored over 6 days via flow cytometry. Coculture with 1D9-BBz T cells did not meaningfully reduce target T-cell viability. **h.** Live-cell flow cytometry of cell-surface P2RX5 levels on 1D9-BBz T cells following 7 days of co-culture with MM.1s cells at 1:8 E:T. T cells from 3 independent *FL/FL* donors.

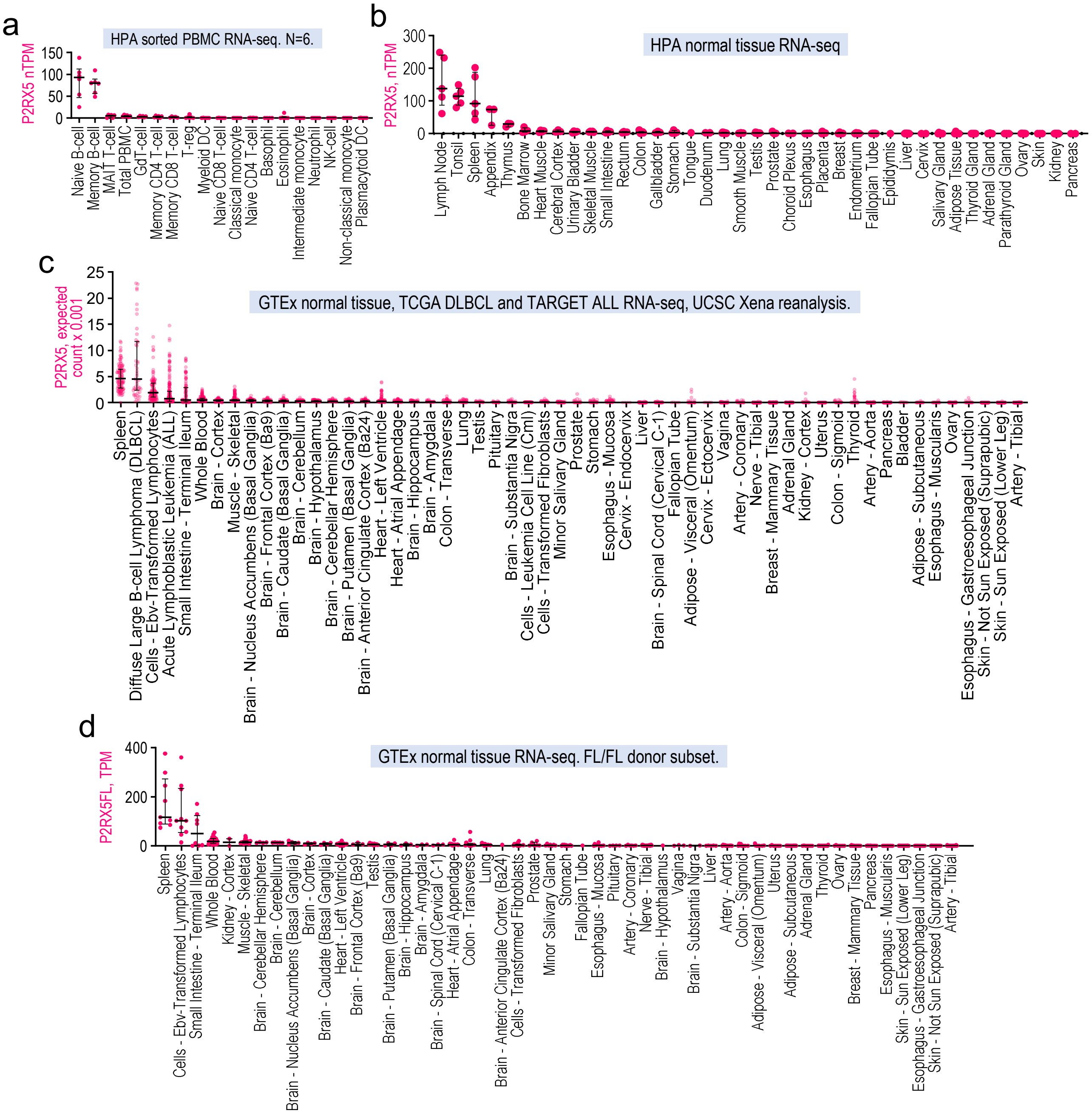


**Fig. 8 | *P2RX5* expression is restricted to sorted B cells and B cell-rich organs such as spleen and lymph nodes. a-d.** Bulk RNA-seq data of P2RX5 transcript levels in: **a**, Human Protein Atlas (HPA) healthy donor cohort sorted peripheral blood mononuclear cells (PBMCs) **b**, HPA cohort normal tissues (reported as nTPM or normalized Transcripts Per Kilobase Million); **c**, TCGA DLBCL and TARGET ALL patient samples as well as GTEx normal tissues (reported as RSEM expected counts, from the UCSC Xena portal); as well as **d**, GTEx normal tissue samples which are homozygous for P2RX5FL (FL/FL), based on our reanalysis of GTEx RNA-seq data (reported as TPM) .

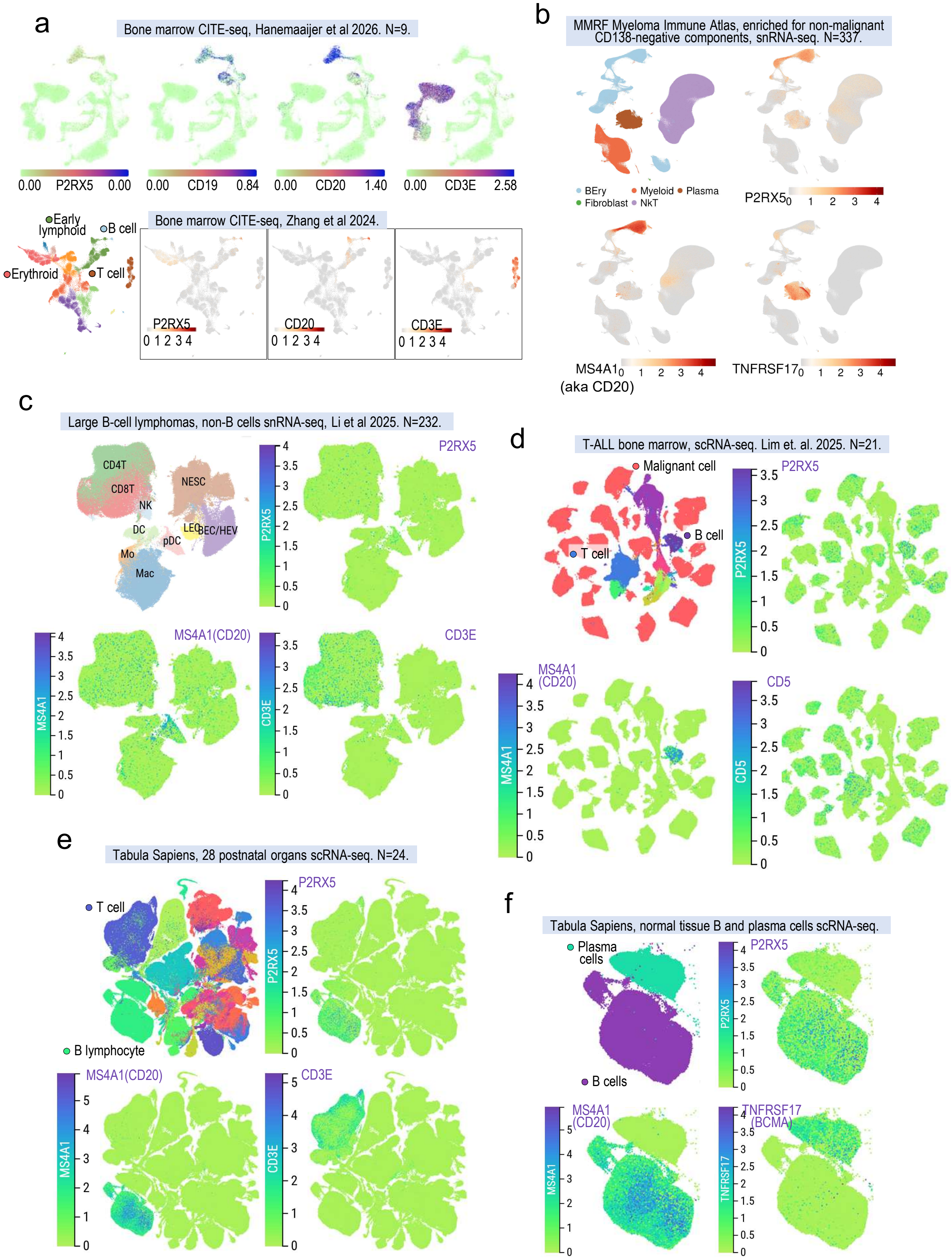


**Fig. 9 | Among non-malignant cells, only B cells and to a lesser extend, plasma cells, upregulate P2RX5 mRNA.** Single cell transcriptomic datasets of P2RX5 expression in: healthy donor bone marrows (**a**); the bone marrow cells of patients with MM (**b**), large B-cell lymphomas (**c**) and T-ALL (**d**); as well as 28 postnatal organs from the Tabula Sapiens v2 portal (**e, f**). Also shown for comparison are the MS4A1(aka CD20) and CD19 B-cell markers, as well as the CD3E T-cell marker. UMAP projections are colored according to the expression level of the indicated gene.

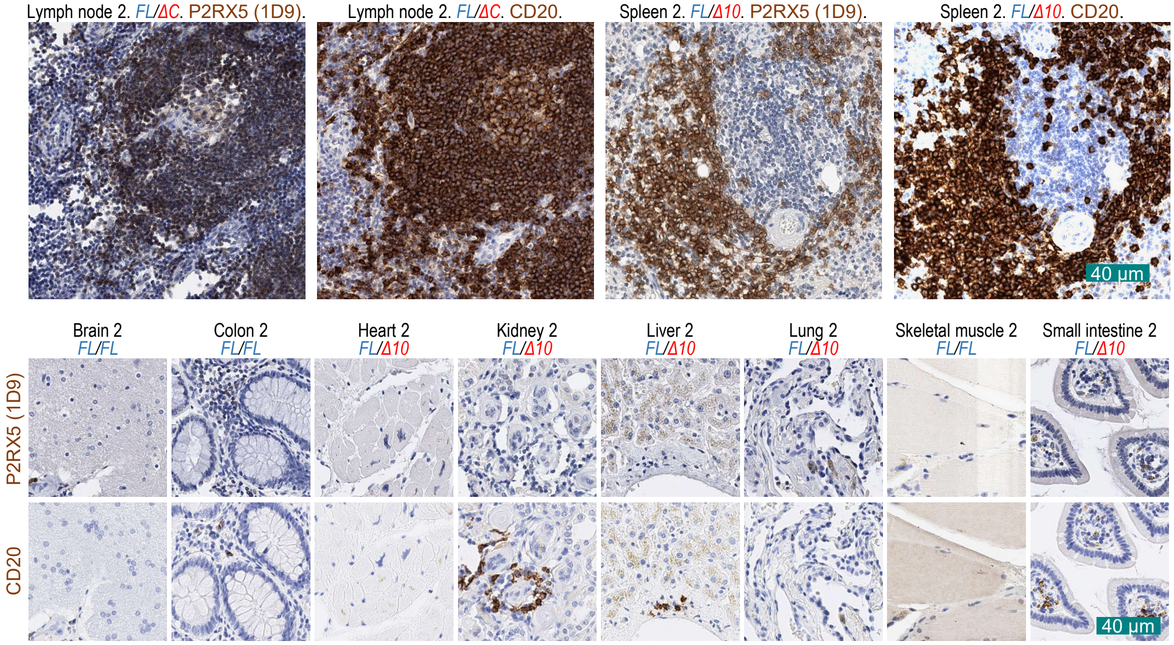


**Fig. 10 | B cell–restricted expression of *P2RX5FL* in normal, postnatal tissues.** Representative IHC images of normal tissue from donors expressing P2RX5 FL/FL or FL/Δ10 or FL/ΔC. FFPE sections were stained for the expression of either P2RX5 (via 1D9 antibody) or CD20 using DAB substrate (brown), with hematoxylin nuclear counterstaining (blue). This is a 2$^{nd}$ independent set of tissue donors from those found in main Fig. 6f.
